# Material aging causes centrosome weakening and disassembly during mitotic exit

**DOI:** 10.1101/866434

**Authors:** Matthäus Mittasch, Vanna M. Tran, Manolo U. Rios, Anatol W. Fritsch, Stephen J. Enos, Beatriz Ferreira Gomes, Alec Bond, Moritz Kreysing, Jeffrey B. Woodruff

## Abstract

Centrosomes must resist microtubule-mediated forces for mitotic chromosome segregation. During mitotic exit, however, centrosomes are deformed and fractured by those same forces, which is a key step in centrosome disassembly. How the functional material properties of centrosomes change throughout the cell cycle, and how they are molecularly tuned remain unknown. Here, we used optically-induced flow perturbations to determine the molecular basis of centrosome strength and ductility in *C. elegans* embryos. We found that both properties declined sharply at anaphase onset, long before natural disassembly. This mechanical transition required PP2A phosphatase and correlated with inactivation of PLK-1 (Polo Kinase) and SPD-2 (Cep192). *In vitro*, PLK-1 and SPD-2 directly protected centrosome scaffolds from force-induced disassembly. Our results suggest that, prior to anaphase, PLK-1 and SPD-2 confer strength and ductility to the centrosome scaffold so that it can resist microtubule-pulling forces. In anaphase, centrosomes lose PLK-1 and SPD-2 and transition to a weak, brittle state that enables force-mediated centrosome disassembly.

## INTRODUCTION

Centrosomes nucleate and anchor microtubules that comprise the mitotic spindle, which segregates chromosomes during somatic cell division. Centrosomes are micron-scale, membrane-less organelles containing a structured centriole pair surrounded by an amorphous protein mass called pericentriolar material (PCM). PCM carries out most of the functions of a centrosome, including directing cell polarity, cell migration, and chromosomal segregation (Conduit et al., 2015; Woodruff et al., 2014)

For chromosome segregation, centrosomes must bear microtubule-dependent loads that create tensile stresses. Motor proteins anchored at the plasma membrane attach to and walk along astral microtubules extending from centrosomes. These spatially-fixed motors thus generate cortically-directed pulling forces on centrosomes, and the balance of those forces determines the ultimate position of the mitotic spindle (Colombo et al., 2003; Gonczy et al., 2001; Grill et al., 2001; McNally, 2013; Nguyen-Ngoc et al., 2007). During this time, centrosomes maintain a compact, spherical shape. However, once chromosome segregation is complete and the cell exits mitosis, centrosomes are deformed and fractured by the same microtubule-mediated forces, which is a pronounced event during centrosome disassembly (Megraw et al., 2002; Severson and Bowerman, 2003). How the cell regulates the structural and material integrity of centrosomes is unclear. One possibility is that an increase in cortical forces during mitotic exit induces centrosome disassembly. In *C. elegans* embryos, the magnitude of microtubule-mediated pulling forces does increase during the metaphase-anaphase transition. Yet, the same increase in pulling forces also occurs in metaphase-arrested embryos without leading to centrosome deformation or fracture (Labbe et al., 2004). Furthermore, artificially increasing pulling forces via *csnk-1* RNAi does not cause premature centrosome disassembly (Magescas et al., 2019; Panbianco et al., 2008). These studies suggest that induction of centrosome deformation and fracture during mitotic exit cannot be sufficiently explained by increased microtubule-mediated forces. An alternative hypothesis is that centrosome mechanical properties significantly change to permit force-driven fracture and dispersal during mitotic exit.

PCM provides most of the mass and microtubule nucleation capacity of a centrosome, and it is widely believed to be responsible for bearing microtubule-mediated forces. PCM is dynamic and expands in size and complexity as cells prepare for mitosis. Self-assembly of coiled-coil proteins, such as Cdk5Rap2 (vertebrates), Centrosomin (*D. melanogaster*) and SPD-5 (*C. elegans*), creates the underlying structural scaffold of PCM which then recruits “client” proteins that nucleate and regulate microtubules (Conduit et al., 2010; Conduit et al., 2014a; Fong et al., 2008; Hamill et al., 2002; Woodruff et al., 2017; Woodruff et al., 2015). Formation of such micron-scale scaffolds requires additional regulatory clients like Polo Kinase, Aurora A Kinase, and SPD-2/Cep192 (Conduit et al., 2014a; Conduit et al., 2014b; Gomez-Ferreria et al., 2007; Hamill et al., 2002; Hannak et al., 2001; Haren et al., 2009; Lee and Rhee, 2011; Pelletier et al., 2004; Zhu et al., 2008). PCM disassembles at the end of each cell cycle, but the mechanism is not well understood. While this process involves microtubule-mediated PCM fracture and reversal of Polo Kinase phosphorylation (Enos et al., 2018; Magescas et al., 2019; Pimenta-Marques et al., 2016), it remains unclear how PCM fracture is initiated, which key molecular targets are de-phosphorylated, if these activities are linked, and how dynamic material changes might contribute to the disassembly process.

A material’s load-bearing capacity is determined by its ability to resist permanent deformation and fracture upon stress. In materials science, these properties are described as “strength” and “ductility”, respectively. Strength is achieved through high affinity bonding and serves to maintain the material’s shape but can sometimes sacrifice flexibility. Ductility is achieved through breakage and reformation of sacrificial bonds or localized neighbor exchange, which dissipates energy over time but sacrifices the material’s shape. For example, glass requires large forces to deform, but it cannot deform much before shattering; thus, glass has high strength and low ductility. On the other hand, chewing gum is easily deformed, and it will stretch to great lengths before breaking; thus, gum has low strength and high ductility. Materials with the highest load-bearing capacity are those that combine strength and ductility, such as rubbers, polyampholyte gels, and high-entropy alloys (George et al., 2019; Sun et al., 2013). Over time, these properties can change via chemical or physical modifications, which is referred to as “material aging”.

For the centrosome, it remains unexplored how molecular-level interactions between PCM proteins generate meso-level mechanical properties like strength and ductility and how these properties change with time. The non-covalent interactions between PCM scaffold proteins, as well as cross-linking of scaffold molecules by PCM clients, could all contribute. In general, characterizing the mechanical properties of living soft matter—such as cells, organelles, and protein assemblies—has been challenging due to their size (sub-micrometer scale) and low abundance (sub-milligram scale). Techniques like atomic force microscopy and optical trapping can be useful in this respect, but they are limited to easily accessible samples, like the outer membrane of cultured cells and reconstituted protein complexes. As a complementary method to actively probe mechanical properties in cells with limited accessibility, we previously used focused light-induced cytoplasmic streaming (FLUCS)(Mittasch et al., 2018). Specifically, we showed how FLUCS can reveal robust power-law-rheology signatures within the cell cytoplasm, to distinguish between fluid and gel-like states. Fluids undergo unconstrained motion proportional to stimulus time (i.e., they flow), while solids and gels undergo only limited deformations, which stall after small amounts of time due to their intrinsic elastic constraints. As FLUCS functions via thermoviscous flows, which develop independent of the absolute viscosity of the fluid (Weinert et al., 2008), FLUCS is particularly suited to distinguish between highly viscous phases and elastic phases. Such a distinction could not be achieved by passive microrheology, as these two states would exhibit the same fingerprint of reduced motion.

The *C. elegans* embryo is an ideal system to dissect the molecular determinants of PCM load-bearing capacity. First, *C. elegans* has a limited core set of proteins needed for rapid PCM assembly and disassembly, most of which are conserved across eukaryotes: PLK-1 (Polo Kinase homolog), SPD-2 (Cep192 homolog), SPD-5 (functional homolog of Centrosomin and Cdk5Rap2), and LET-92^SUR-6^ (PP2A^B55^*^α^* phosphatase homolog) (Decker et al., 2011; Enos et al., 2018; Hamill et al., 2002; Kemp et al., 2004; Magescas et al., 2019; Pelletier et al., 2004; Schlaitz et al., 2007). Second, it is possible to reconstitute PCM assembly and microtubule nucleation *in vitro* using purified *C. elegans* proteins (Woodruff et al., 2017; Woodruff et al., 2015). These experiments previously revealed that PCM forms via self-assembly of SPD-5 into spherical, micron-scale scaffolds that recruit PCM client proteins. SPD-2 and PLK-1 enhance SPD-5 self-assembly, while PP2A^SUR-6^ removes PLK-1-derived phosphates and promotes PCM disassembly. However, these experiments did not reveal the mechanical properties of the SPD-5 scaffold nor how they are tuned in a cell-cycle-dependent manner.

Here, we ask 1) how centrosomes undergo dynamic structural changes to withstand high tensile stresses in mitosis but not during mitotic exit, 2) which mechanical properties are associated with these distinct functional states, and 3) which molecular players and logics regulate the transient existence of centrosomes. To answer these questions, we combined genetics and pharmacological intervention with FLUCS to study the mechanical properties of PCM in *C. elegans* embryos. Our results revealed that PCM transitions from a strong, ductile state in metaphase to a weak, brittle state in anaphase. This mechanical transition is promoted by PP2A^SUR-6^ and opposed by PLK-1 and SPD-2. Our data suggest that mitotic PCM is a composite of a stable SPD-5 scaffold and proteins that dynamically reinforce the scaffold, such as PLK-1 and SPD-2. During spindle assembly, accumulation of PLK-1 and SPD-2 render the PCM tough enough to resist microtubule-pulling forces. During mitotic exit, departure of PLK-1 and SPD-2 weakens the PCM scaffold to allow microtubule-mediated fracture and disassembly. Thus, PCM undergoes cell-cycle-regulated material aging that functions to promote its disassembly.

## RESULTS

### FLUCS reveals weakening of the PCM scaffold at anaphase onset in *C. elegans* embryos

To study the molecular determinants of PCM load-bearing capacity, we studied *C. elegans* 1-cell embryos, where growth and disassembly of the PCM scaffold is easily visualized using fluorescently-labeled SPD-5 (mMaple::SPD-5)(Figure 1A). During spindle assembly, the *C. elegans* PCM scaffold is subject to microtubule-mediated pulling forces, but it maintains its spherical shape and structural integrity. However, during telophase, those same pulling forces deform and fracture the PCM scaffold (Figure 1A)(Enos et al., 2018; Severson and Bowerman, 2003). We hypothesized that PCM undergoes an intrinsic mechanical transition from a strong, tough state in metaphase to a weak state in telophase.

**Figure 1.**
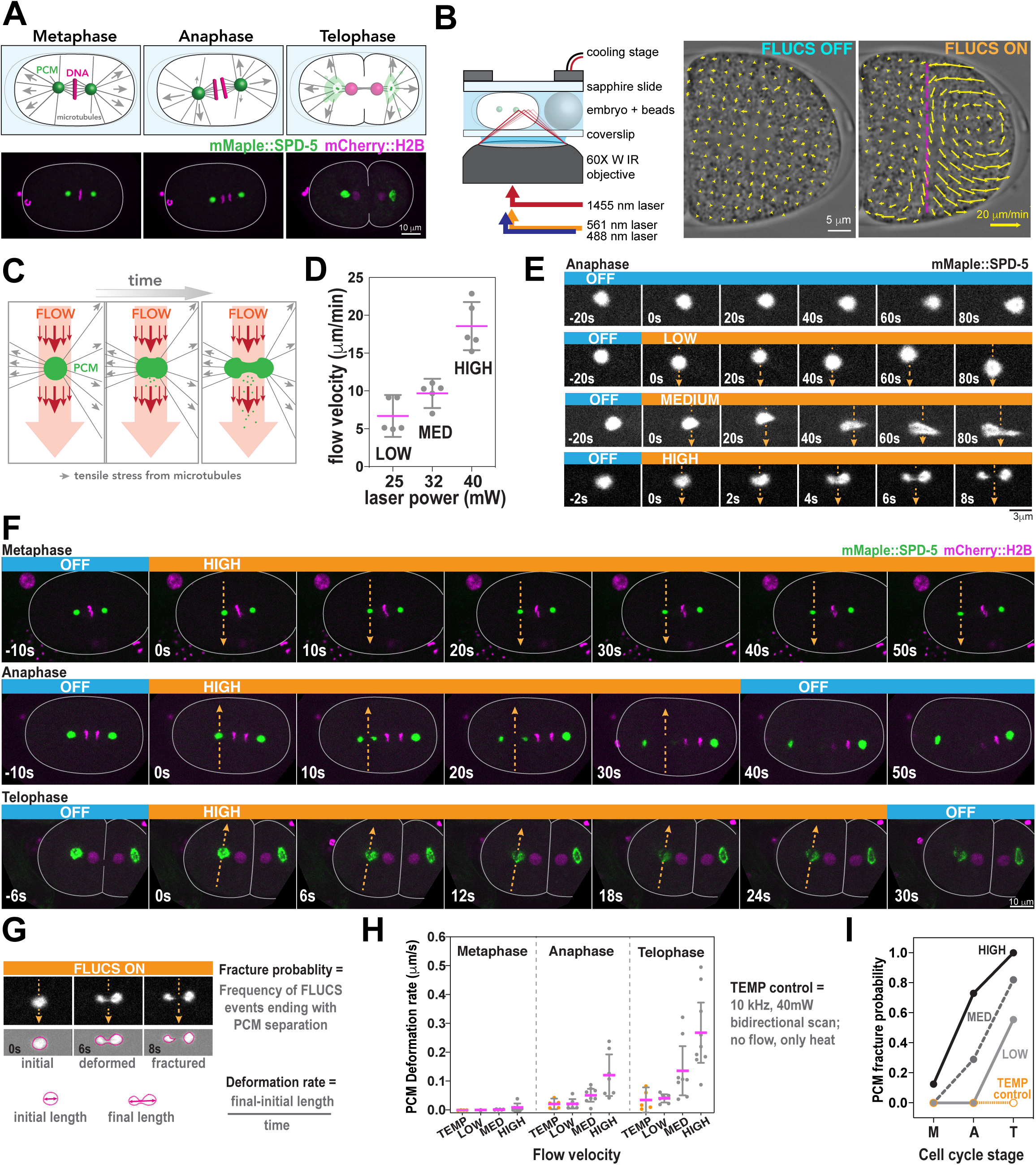
FLUCS reveals changes in PCM deformation resistance and fracture resistance during mitosis in *C. elegans* embryos. A. Diagram of mitotic progression in a one-cell *C. elegans* embryo (top panels). Force generation (arrows) by cortically anchored microtubules aid in chromosome segregation, spindle positioning, and PCM disassembly during telophase. Live-cell confocal microscopy images of *C. elegans* embryos expressing a PCM marker (mMaple::SPD-5) and DNA marker (mCherry::HistoneH2B). B. Diagram of the FLUCS microscope setup (left) and generation of intracellular flows after unidirectional scanning of a 1455 nm laser at 1500 Hz. Scan path is represented by the magenta line. The magnitudes of local flow velocities are reflected by arrow size. C. Using FLUCS, flows (red arrows) are generated in the cytoplasm and pass through the PCM (green ball). Microtubule-derived pulling forces (grey arrows) also exert tensile stresses on PCM. D. Tuning the power of the 1455 nm laser (25, 32, and 40 mW) generates three tiers of flow velocity (LOW, MEDIUM, and HIGH). Individual data points are plotted with mean +/− 95% CI; n = 5 measurements per condition. E. Time lapse images of PCM morphology in anaphase after application of no flow (OFF) or low, medium, and high flow. Orange heading boxes indicate when flow occurs. Arrows indicate flow path and direction. Blue heading boxes indicate when flow is turned off. F. PCM was subjected to high-flow FLUCS during metaphase, anaphase, and telophase. G. For each FLUCS experiment, we measured the change in PCM length over time (Deformation rate) and the frequency of complete separations in PCM (Fracture probability). H. PCM deformation rates were measured in metaphase, anaphase, and telophase using low, medium, and high flow. Individual data points are plotted with mean +/− 95% CI; n = 7,6,7 (metaphase; LOW, MED, HIGH flow), n = 7,8,7 (anaphase), n = 7,8,9 (telophase) measurements per condition. 10 kHz bidirectional scanning of the 1455 nm laser using 40 mW power, generates heat without producing flows (TEMP control; n = 4,5,5 (metaphase, anaphase, telophase)). For high flow, differences are statistically significant using a one-way ANOVA followed by a Tukey’s multiple comparison test (metaphase vs. anaphase, p= 0.04; metaphase vs. telophase, p = 0.0001). I. PCM fracture probabilities from experiments in (H). Sample numbers are the same as in (H).

We probed PCM mechanical properties using FLUCS-generated flows complemented with fluorescent imaging. Specifically, we equipped a spinning disk confocal microscope with a laser control unit (wavelength = 1455 nm) that creates precise, sub-millisecond thermal manipulations (Figure 1B). Unidirectional scans with this laser at 1500 Hz creates travelling temperature fields that are sufficient to induce flows in a viscous medium, including embryonic cytoplasm (Mittasch et al., 2018)(see Methods). A Peltier-cooled stage insert dissipates excess heat to keep the sample within its physiological temperature range.

We applied FLUCS to *C. elegans* embryos expressing mMaple::SPD-5 and mCherry-labeled histones (mCherry::H2B). For each experiment, we induced flows crossing the cytoplasm and continuing through the middle of a centrosome, which should apply stress to the PCM scaffold orthogonal to microtubule-derived tensile stresses (Figure 1C). Based on previous experiments (Mittasch et al., 2018), we predicted that cytoplasmic shear flows should weaken the PCM by the relative displacement of scaffold proteins and create a virtual “notch” in the flow path. We used three different amplitudes for the scanning infrared laser (25, 32, 40 mW) to generate cytoplasmic flow velocities ranging from 5-20 μm/min (Figure 1D; Movie S1); these flows scaled quadratically with laser power (R^2^= 0.97), as one would expect for a predominantly viscous medium (Figure S1A). Simultaneously, we cooled embryos to 17°C, such that the embryo cytoplasm never exceeded 23°C during laser scanning (*C. elegans* embryos develop properly at any temperature between 16°C and 25°C)(Begasse et al., 2015).

As shown in Figure 1E, FLUCS deformed and fractured mature PCM in anaphase embryos. We also detected a slight increase in cytoplasmic fluorescence surrounding the PCM during FLUCS, indicating that flows can dislodge SPD-5 from the PCM, as predicted (Figure S1B). FLUCS-induced PCM deformation differed starkly between cell cycle stages. FLUCS was not able to visibly deform PCM during metaphase or prior, even though flows were strong enough to detach the centrosome from the spindle and move it toward the cell cortex or out of the focal plane (Figure 1D and S1C and Movies S2, S3; see Metaphase and Prometaphase). On the contrary, FLUCS deformed and eventually fractured PCM during anaphase and telophase (Figure 1E,F; Movie S4,S5); in these experiments, the untreated centrosome remained intact.

We then quantified 1) the deformation rate, defined by the rate of PCM length change orthogonal to the flow direction, and 2) the fracture probability, defined by the chance that PCM segments detach completely after FLUCS (Figure 1G; see methods). The PCM deformation rate and fracture probability increased with increasing flow velocity and progression through mitosis (Figure 1 H,I). To pinpoint when PCM becomes susceptible to FLUCS-induced deformation, we continuously applied FLUCS to centrosomes starting in metaphase continuing into anaphase. PCM remained spherical and intact during metaphase, but then fractured immediately after anaphase onset, as marked by chromosome segregation (Figure S1D; Movie S6). Our results suggest that PCM resistance to deformation and fracture is high during metaphase, then declines at anaphase onset, ∼150s prior to full PCM disassembly in telophase. We refer to this change in PCM mechanical properties hereon as the “PCM weakening transition”.

Since the generation of FLUCS is accompanied by local temperature gradients, we tested if temperature alone affects PCM structure. Bidirectional scanning at 10 kHz creates local temperature gradients without flow, and these conditions did not cause significant PCM deformation or fracture (Figure 1H,I and S1E). Thus, centrosome perturbation during FLUCS is primarily due to the flows and not temperature gradients *per se*. Furthermore, embryos developed properly after cessation of FLUCS (Figure S1F)(Mittasch et al., 2018). We conclude that this established method can be used to study organelles inside a living cell.

### PCM weakens in anaphase independent of cortical force generation

We next wondered if our FLUCS results could be explained by cell-cycle-dependent changes in PCM mechanical properties or changes in cortical force generation. During the metaphase-anaphase transition in early *C. elegans* embryos, cortical pulling forces increase to induce transverse oscillations and posterior positioning of the mitotic spindle (Pecreaux et al., 2006). Depleting the proteins GPR-1 and GPR-2 (*gpr-1/2(RNAi)*) significantly reduces cortical microtubule-pulling forces and prevents spindle displacement, spindle oscillation, and PCM deformation and fracture (Colombo et al., 2003; Enos et al., 2018; Grill et al., 2003; Magescas et al., 2019; Pecreaux et al., 2006). Thus, we performed FLUCS in *gpr-1/2(RNAi)* embryos, where we expect only residual cortical forces that remain relatively constant throughout mitosis.

In both wild-type and *gpr-1/2(RNAi)* embryos, FLUCS deformed PCM in anaphase and telophase, but not in metaphase. However, in *gpr-1/2(RNAi)* embryos, FLUCS-induced PCM deformation rates were slower (Figure 2 A,B; Figure S2A), and FLUCS caused fracturing only in telophase (Figure 2C). These results suggest that PCM mechanical properties change during mitotic exit, but that cortical pulling forces are required to disperse and fracture the pre-weakened PCM scaffold. To further test this idea, we applied 10 μg/ml nocodazole to depolymerize microtubules and performed high-flow FLUCS. Under these conditions, FLUCS did not visibly affect PCM during metaphase, but did dislodge SPD-5 protein from the PCM during telophase (Figure 2D); we did not observe stretching or clean fracture of the PCM during any cell cycle stage. These results indicate that 1) the interactions between SPD-5 scaffold molecules weaken independent of microtubule-pulling forces during mitotic exit, and 2) microtubule-pulling forces are required for stretching and fracture of the PCM during mitotic exit. To test if microtubule-mediated pulling forces could be sufficient to deform PCM already in metaphase, we depleted a negative regulator of GPR-1/2, casein kinase 1 gamma (*csnk-1(RNAi)),* which is reported to increase cortical pulling forces ∼1.5-fold (Panbianco et al., 2008). We did not observe premature deformation or fracture of PCM under these conditions, even though the spindle rocked violently in metaphase (Figure S2B). We conclude that an intrinsic mechanical change in the SPD-5 scaffold is the main driver of PCM weakening during anaphase entry.

**Figure 2.**
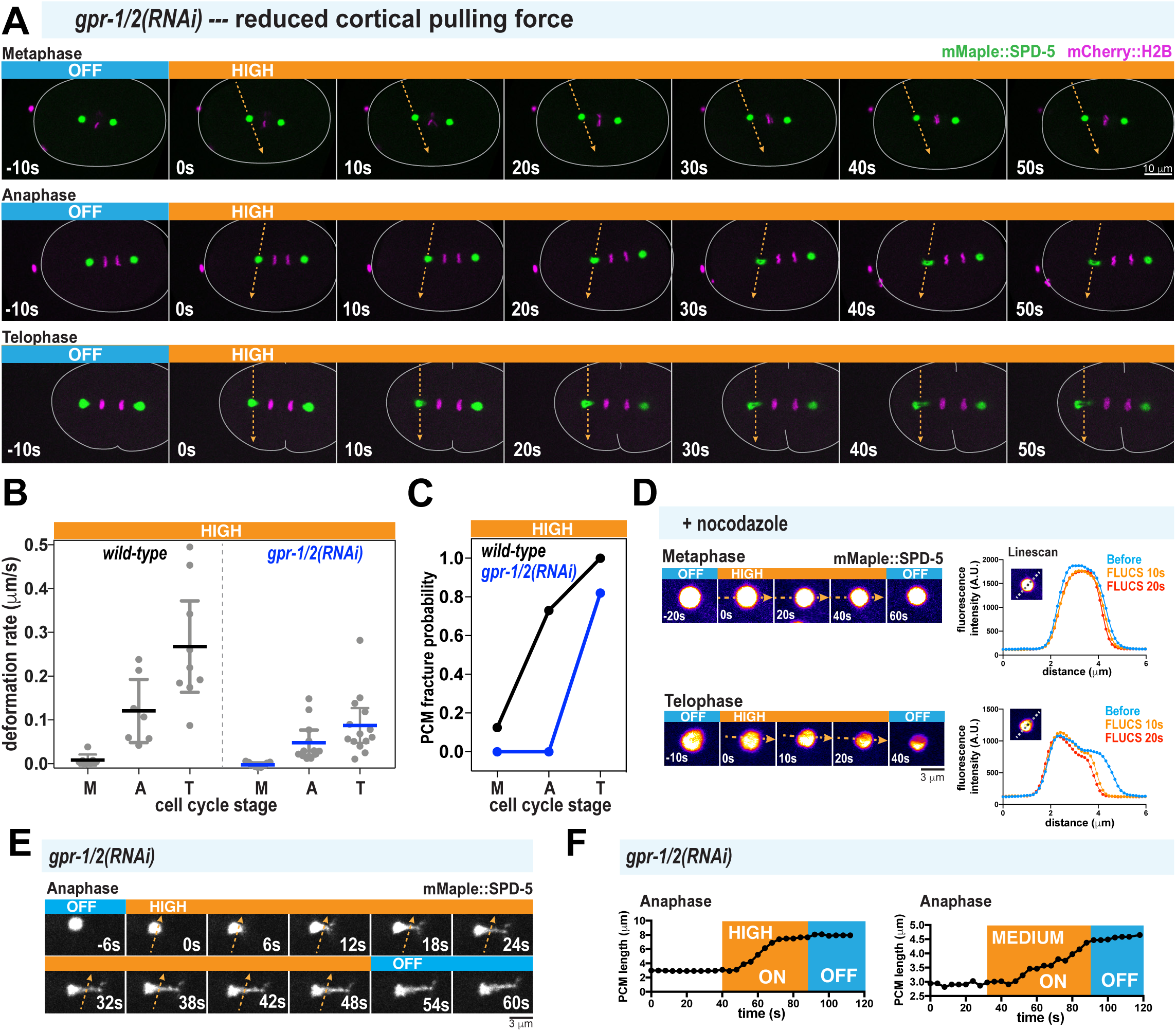
PCM undergoes structural weakening during anaphase, independent of cortical pulling forces. A. Time-lapse images of *gpr-1/2(RNAi)* embryos treated with 40 mW FLUCS (high flow). B. PCM deformation rates in metaphase (M), anaphase (A), and telophase (T) using high flow in wild-type and *gpr-1/2(RNAi)* embryos. Wild-type data are from experiments in Figure 1. Individual data points are plotted with mean +/− 95% CI; n = 7,7,9 (wild-type; metaphase, anaphase, telophase) and n = 10,12,14 (*gpr-1/2(RNAi)*). C. PCM fracture probabilities from experiments in (B). D. Permeabilized embryos were treated in metaphase or telophase with 10 μg/ml nocodazole, then subjected to high-flow FLUCS. Representative images are shown on the left, line scans (dotted line in the inset) of fluorescence intensity before and after FLUCS are on the right. E. Zoomed in time-lapse images of PCM deformation under high-flow FLUCS in a *gpr-1/2(RNAi)* embryo. F. Plots comparing PCM length on the long axis orthogonal to flow direction over time. Both high flow and medium flow induce PCM deformation.

We next examined the viscoelastic properties of PCM during anaphase by measuring the time-dependent deformation of PCM during FLUCS and relaxation after FLUCS was turned off. We used *gpr-1/2(RNAi)* embryos to allow residual pulling forces but prevent spindle oscillations, which could complicate our analysis. We observed that continuous application of medium and high-flow FLUCS in *gpr-1/2(RNAi)* embryos caused time-dependent strain of the PCM scaffold (Figure 2E). Thus, anaphase PCM is ductile and can experience micron-scale structural rearrangements without complete fracture during stress. Such behavior is seen in viscous materials. When we turned off FLUCS, PCM remained in its strained, elongated state, indicating the absence of a dominant elastic element strong enough to return the PCM to its original shape (Figure 2 E,F).

Overall, our FLUCS experiments suggest that the PCM structurally weakens after metaphase. This weakening transition would presumably facilitate PCM disassembly by enabling microtubule-dependent pulling forces to fracture and disperse the PCM scaffold in telophase.

### PCM undergoes stepwise compositional changes following anaphase onset

We next investigated the molecular mechanism underlying the PCM weakening transition, in particular, identifying the specific players that determine the dynamic regulation of PCM strength and ductility. PCM is a heterogeneous assembly of proteins required for its assembly and function (Figure 3A). In particular, two critical regulatory proteins, PLK-1 (Polo-like Kinase) and SPD-2 (Cep192 homolog), interact with the scaffold protein SPD-5 and enhance its self-assembly into supramolecular structures (Cabral et al., 2019; Decker et al., 2011; Woodruff et al., 2017; Woodruff et al., 2015). PLK-1 and SPD-2, as well as other PCM-localized client proteins, might also reinforce the mature SPD-5 scaffold. On the other hand, loss or inactivation of these proteins could weaken the PCM scaffold.

**Figure 3.**
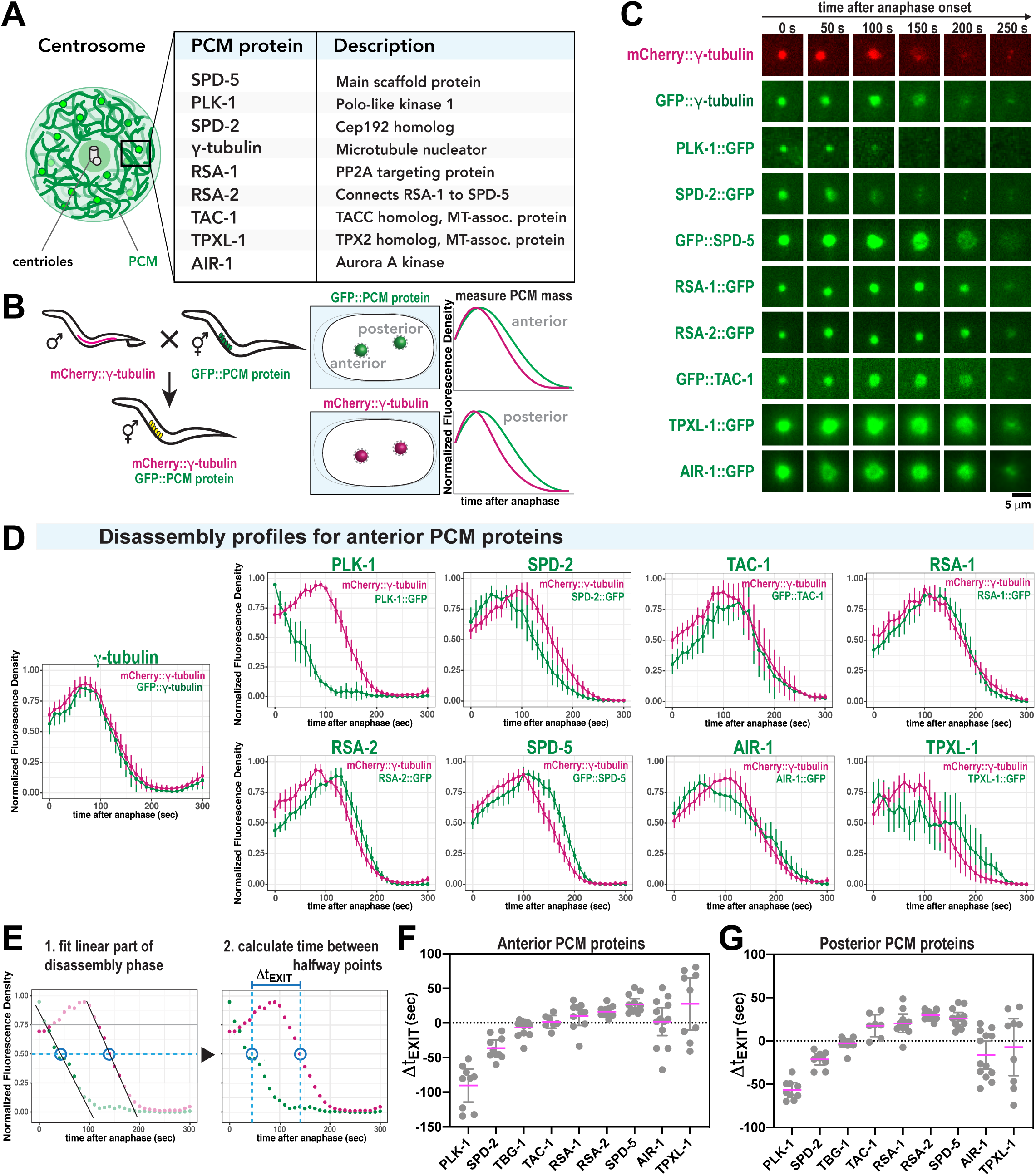
Discrete changes in PCM composition correlate with the PCM weakening transition in anaphase. A. Diagram of *C. elegans* centrosome architecture and composition. B. Worm lines were generated that express mCherry-labeled *γ*-tubulin (as a standard) and 9 different GFP-labeled PCM proteins (left panels). Fluorescence intensity at the PCM was measured over time (right panels). C. Example images from dual-color, time-lapse recording of PCM disassembly in 9 different embryo lines described in (B). D. Quantification of the experiments in B-C. For each strain, the plots represent the normalized integrated fluorescence density of PCM-localized mCherry-tagged *γ*-tubulin compared to the GFP-tagged protein from anaphase onward. Anaphase was indicated by spindle rocking. Shown are the analyses for anterior-localized centrosomes. Data were normalized to the maxima for each individual curve, then these curves were averaged (mean +/− 95% C.I. n = 13 (*γ*-tubulin), 9 (PLK-1), 10 (SPD-2), 12 (SPD-5), 14, (RSA-1), 12 (RSA-2), 7 (TAC-1), 10 (TPXL-1), 13 (AIR-1)). E. The order of PCM protein departure was determined by calculating the time lag between halfway points of PCM protein departure per strain (*Δ*t_EXIT_). Halfway points were determined by fitting each curve during the window of linear departure. F. Departure time lag (*Δ*t_EXIT_) of GFP-labeled PCM proteins relative to mCherry::*γ*-tubulin. A negative value indicates that the GFP-labeled protein departed before *γ*-tubulin. A positive value indicates that the GFP-labeled protein departed after *γ*-tubulin. Results for anterior centrosomes are shown (mean +/− 95% C.I.; sample number is the same as in (D)). Statistical analyses are shown in Table S3. G. Departure time lag (*Δ*t_EXIT_) of posterior-localized PCM proteins relative to mCherry:: *γ*-tubulin. Sample number is the same as in (D). Statistical analyses are shown in Table S4.

To analyze PCM composition changes during anaphase, we visualized 9 different GFP-labeled PCM proteins relative to a standard PCM marker, mCherry::*γ*-tubulin. We then measured the integrated density of PCM-localized mCherry and GFP signals during mitosis (Figure 3B). The curves in Figure 3D and Figure S3 represent averages for >10 experiments (mean +/− 95% CI). mCherry::*γ*-tubulin signal peaked ∼75-100s after anaphase onset, then declined, indicating its departure from PCM. GFP::*γ*-tubulin behaved similarly, as expected (Figure 3C,D and Figure S3). PLK-1 signal decreased immediately after anaphase onset and was no longer detectable ∼100s later. SPD-2 also departed from the PCM prior to *γ*-tubulin. However, the main scaffold protein SPD-5 departed afterward. All other proteins departed coincidentally with *γ*-tubulin or soon afterward. TPXL-1 and AIR-1 departed in a biphasic manner: an initial loss of signal occurred prior to *γ*-tubulin departure, then a second phase occurred after *γ*-tubulin departure. To compare departure kinetics across all experiments, we determined the halfway point of disassembly for each individual GFP and mCherry curve per experiment, then calculated the time differential between halfway points (*Δ*t_EXIT_; Figure 3E). The results for anterior and posterior PCM proteins are summarized in Figure 3F and 3G, respectively. A negative *Δ*t_EXIT_ value indicates GFP::PCM protein departure before *γ*-tubulin, and a positive value indicates departure after *γ*-tubulin. Our results reveal that PCM composition changes in a stepwise manner during anaphase: PLK-1 departs first, followed by SPD-2, *γ*-tubulin, TAC-1, and finally SPD-5 and proteins that form tight complexes with SPD-5 (RSA-1, RSA-2). TPXL-1 and AIR-1 were more variable in their departure, possibly because they localize both to PCM and microtubules that remain after disassembly of the PCM scaffold (Hannak et al., 2001; Ozlu et al., 2005).

### Polo Kinase and SPD-2 reinforce the PCM scaffold by increasing its strength and ductility

PLK-1 and SPD-2 are the first proteins to depart the PCM during anaphase, when the PCM begins to weaken. Thus, we hypothesized that PLK-1 and SPD-2 normally reinforce the PCM to achieve full strength and stability in metaphase. If this idea is correct, then acute inhibition of PLK-1 phosphorylation or SPD-2 in metaphase might prematurely weaken the PCM, accelerate its disassembly, or reveal hidden material states not previously visible.

For acute inhibition of PLK-1, we treated semi-permeable embryos (via *perm-1(RNAi)*) with 10 μM Polo Kinase inhibitor BI-2536 in prometaphase (Carvalho et al., 2011). After 2 minutes in drug solution, we applied low, medium, and high-flow FLUCS to centrosomes (Figure 4A-C). Under these conditions, low and medium-flow FLUCS deformed metaphase PCM in BI-2536-treated embryos, in contrast to wild-type embryos (Figure 4B); in both cases, PCM fracture did not occur. Under high-flow FLUCS, BI2536 treatment increased PCM deformation rate as much as ∼11-fold with fracture occurring only in a minority of the cases (30%) (Figure 4A,C). The fact that PLK-1 inhibition enabled PCM to be deformed easily without necessarily fracturing suggests that PLK-1 mostly determines PCM strength. This experiment also reveals that wild-type PCM is ductile during metaphase; this could not be observed previously because the deformation resistance of wild-type PCM was too high. BI-2536 treatment also caused premature disassembly of the SPD-5 scaffold in metaphase-arrested embryos, consistent with previous findings (Figure 4D)(Cabral et al., 2019). Our results show that continuous PLK-1 activity is needed for PCM to achieve full strength and maintain integrity until chromosome are separated.

**Figure 4.**
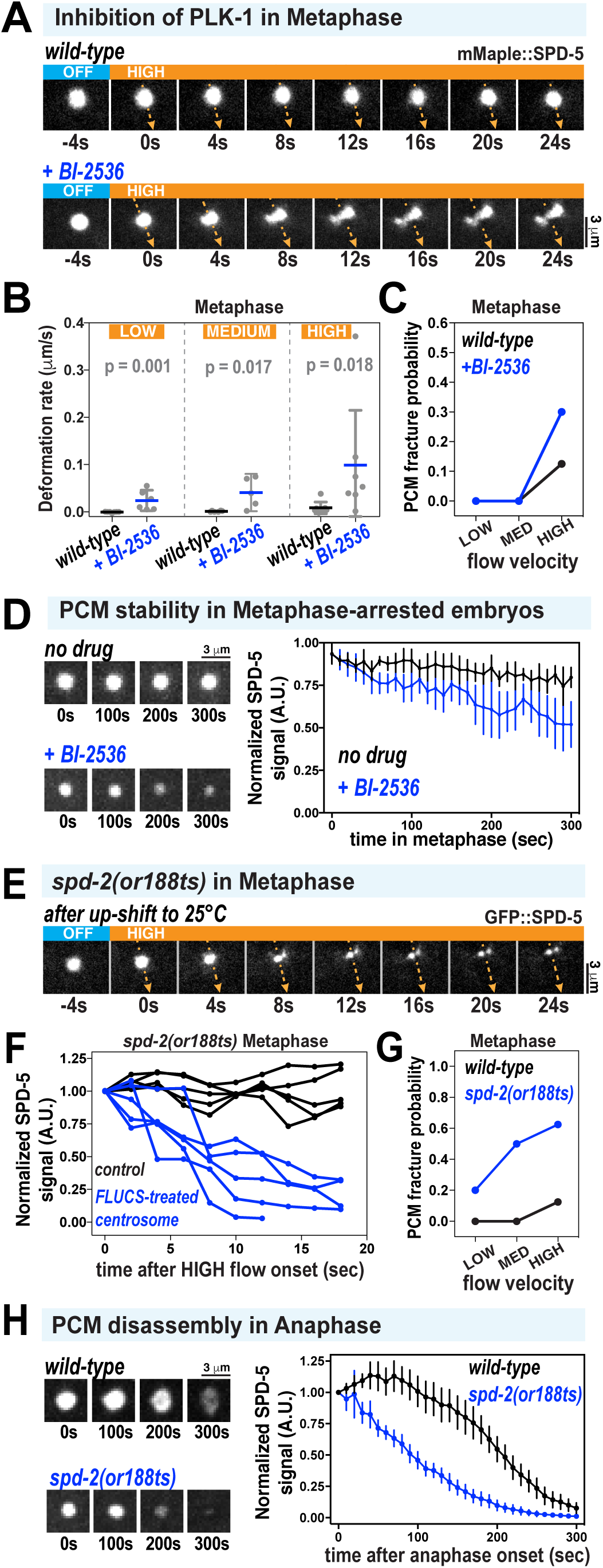
Acute inhibition of PLK-1 and SPD-2 induces premature weakening and disassembly of the PCM scaffold. A. PCM was subjected to high-flow FLUCS during metaphase in wild-type embryos or permeabilized embryos treated with 10 μM BI-2536 (inhibitor of Polo Kinases). Permeabilized embryos behaved as wild-type embryos during the first cell division (see methods; Carvalho 2011). B. PCM deformation rates in metaphase using low, medium, and high flow in wild-type and BI-2536-treated embryos. Wild-type data are from experiments in Figure Individual data points are plotted showing mean +/− 95% CI; n = 6-7 (wild-type) and n = 5-7 (BI-2536-treated). P-values were calculated using a Mann-Whitney test. C. PCM fracture probabilities from experiments in (B). D. Permeabilized embryos were arrested in metaphase using 10 μM MG-132, then treated with 0.1% ethanol (no drug) or 10 μM BI-2536. Data are plotted as normalized lines representing mean +/− 95% CI; n = 8 (no drug) and n = 10 (BI-2536-treated). E. Embryos expressing GFP::SPD-5 and a temperature-sensitive version of SPD-2 (*spd-2(or188ts)*) were allowed to assemble centrosomes at the permissive temperature (16°C), upshifted to the non-permissive temperature (25°C) for 1 min during prometaphase, then subjected to high-flow FLUCS during metaphase. F. For each experiment in *spd-2(or188ts)* embryos, one centrosome was subjected to FLUCS and the other left alone (control). Integrated fluorescent intensities of the SPD-5 signal were tracked over time, then normalized to the starting value. Each curve represents a single experiment. G. PCM fracture probabilities using low, medium, and high flow. Wild-type data are reproduced from Figure 1. n = 7,7,8 (wild-type) and 5,6,8 (*spd-2(or188ts)*). H. Embryos were upshifted from 16°C to 23°C during metaphase, then imaged during anaphase. Data show integrated fluorescence densities of PCM-localized signal, plotted as normalized lines representing mean +/− 95% CI; n = 24 centrosomes in both wild-type and *spd-2(or188ts)* embryos.

Next, we analyzed embryos expressing a temperature-sensitive version of SPD-2 (*spd-2(or188ts)*)(Kemp et al., 2004). We mounted *spd-2(or188ts) gfp::spd-5* embryos in cold media and maintained the sample at 17°C while imaging until prometaphase, then upshifted the embryos to 25°C for 1 min to inactivate SPD-2^or188ts^. We then lowered the temperature to 17°C to perform FLUCS in metaphase as per usual (Figure 4E)(Note: because of the local heating caused by FLUCS, the treated centrosome remained at ∼23°C throughout the experiments). The absence of fully functional SPD-2 made PCM more susceptible to FLUCS-induced fracture and disintegration at all applied flow velocities (Figure 4E-G). Even in PCM that did not fracture into observable pieces, the GFP::SPD-5 signal decayed after application of FLUCS (Figure 4F). We did not observe this phenotype in wild-type PCM or *spd-2* mutant PCM not treated with FLUCS (Figure 4F). Our interpretation of this data is that SPD-2 is required for PCM ductility and strength. Without SPD-2, PCM becomes brittle and susceptible to fracture and diffusion-driven departure of constituents after modest mechanical agitation. In line with this view, inactivation of SPD-2 caused premature disassembly of the SPD-5 scaffold in early anaphase, even without FLUCS perturbations (Figure 4H). Deformation rates were difficult to measure because rapid fracture and vanishing GFP::SPD-5 signal precluded a flow analysis. We conclude that both Polo Kinase and SPD-2 help PCM achieve maximal strength and ductility to prevent disassembly.

We next used a minimal *in vitro* system to test if PLK-1 and SPD-2 directly affect the mechanical properties of the SPD-5 scaffold. When incubated in a crowded environment (e.g. >4% PEG), purified recombinant SPD-5 assembles into micron-scale condensates that recruit PLK-1, SPD-2, and other PCM proteins (Woodruff et al., 2017). We could not assess SPD-5 condensates using FLUCS because the condensates were propelled quickly away from the flow path (data not shown); thus, our simplified in vitro conditions do not exactly match those found in native cytoplasm. As another way to assess the strength of SPD-5 interactions, we induced disassembly of young RFP-labeled SPD-5 condensates (500 nM SPD-5::RFP; 5 min after formation) through application of pipetting shear forces and dilution, then measured the amount of condensates that survived using fluorescence microscopy (Figure 5A)(note: dilution is required to prevent SPD-5 re-assembly; thus, this assay tests resistance to disassembly only)(Enos et al., 2018). This treatment completely disassembled condensates composed solely of SPD-5 (Figure 5B,C). Addition of constitutively active PLK-1 (PLK-1^CA^; T194D T-loop phospho-mimic) or SPD-2 prevented SPD-5 condensate disassembly, with the combination of the two yielding the greatest protection (Figure 5B,C). Kinase-dead PLK-1 (PLK-1^KD^; K67M mutant) did not promote SPD-5 condensate survival. These results suggest that PLK-1 phosphorylation of SPD-5, along with direct binding of SPD-2, reinforce the interactions between SPD-5 molecules and thus enhance the ability of PCM to resist disassembly.

**Figure 5.**
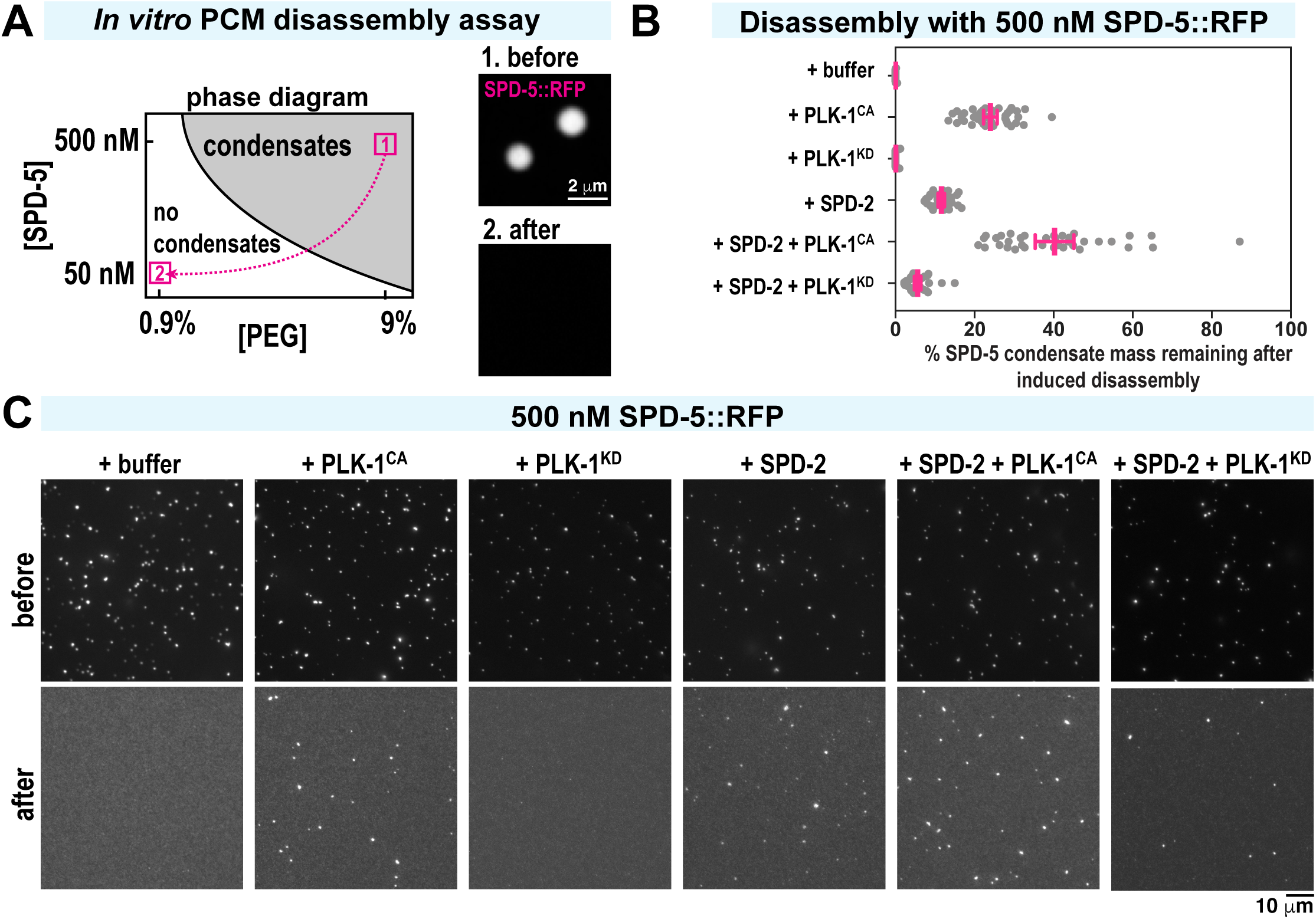
Polo Kinase and SPD-2 protect *in vitro* reconstituted PCM from induced disassembly. A. *In vitro* SPD-5 condensate disassembly experiment. 500 nM SPD-5::TagRFP was incubated in 9% PEG to induce spontaneous formation of micron-scale SPD-5 condensates (1. before). After 5 min, the condensates were pipetted harshly and diluted 1:10 in PEG-free buffer, incubated for 10 min, then imaged (2. after). B. Quantification of total SPD-5 condensate mass per field of view remaining after dilution-induced disassembly. Buffer, 240 nM SPD-2, 500 nM constitutively active PLK-1 (PLK-1^CA^), and/or 500 nM kinase dead PLK-1 (PLK-1^KD^) were added at the beginning. The plot shows total integrated fluorescence density for each field of view (red bars indicate mean +/− 95% C.I.; n >22 images per experiment). C. Representative images from (B) before and after dilution-induced disassembly.

Our *in vitro* and *in vivo* data together suggest that PLK-1 and SPD-2 tune PCM load-bearing capacity by conferring strength and ductility to the SPD-5 scaffold.

### Phosphatase PP2A^SUR-6^ promotes PCM disassembly by compromising scaffold ductility

We next investigated how embryos promote the PCM weakening transition during anaphase. PP2A phosphatase localizes to the PCM and plays multiple roles in centriole biogenesis, spindle assembly, and mitotic exit (Wlodarchak and Xing, 2016). The *C. elegans* homolog of PP2A (LET-92) complexed with the B55*α* regulatory subunit SUR-6 (PP2A^SUR-6^) is required for complete PCM disassembly (Enos et al., 2018; Magescas et al., 2019). We thus tested if PP2A^SUR-6^ drives PCM disassembly by compromising the mechanical properties of PCM.

We treated semi-permeable one-cell embryos with 10 μM PP2A inhibitor (LB-100) in metaphase, then performed high-flow FLUCS in anaphase. Unlike in wild-type embryos, where PCM fractured quickly after high-flow FLUCS, the PCM in LB-100-treated embryos stretched orthogonal to the induced flow but resisted fracture (Figure 6A). PCM deformation velocity was ∼2-fold higher (0.26 vs. 0.12 μm/min), initially suggesting that PCM is easier to deform when PP2A is inhibited (Figure 6B). However, PP2A inhibition also elevated the ductility of PCM 1.5-fold (final length divided by the original length) and lowered the fracture probability >2-fold in all cell cycle stages (Figure 6C-D). In 2/10 anaphase embryos treated with LB-100, PCM stretched as much as 4-fold in length after FLUCS, reaching up to 10 μm while staying connected. PCM was also more resistant to fracture in embryos depleted of the PP2A regulatory subunit SUR-6 (Figure S4A-C). These results suggest that, when PP2A is inhibited, the ductile nature of PCM is preserved throughout anaphase, allowing PCM to absorb more energy overall without fracturing. This is likely achieved through “self-healing”, or the breakage and reformation of weak inter-scaffold interactions. The increase in deformation velocity may then result from ductile PCM becoming easier to stretch as it becomes more extended. On the other hand, wild-type PCM is brittle during anaphase and can only be extended short distances before fracturing. We conclude that PP2A normally functions to eliminate “self-healing” PCM scaffold interactions, thus making PCM brittle during anaphase and susceptible to microtubule-mediated fracture in telophase. Consistent with this conclusion, LB-100 inhibition of PP2A or depletion of its regulatory subunit SUR-6 inhibited SPD-5 scaffold disassembly in telophase (Figure 6E) (Enos et al., 2018). We speculate that PCM may be less porous in this mutant ductile state compared to the wild-type brittle state, which could delay PCM disassembly further by preventing access of additional disassembly machinery and/or delaying the departure of PLK-1 and SPD-2. Consistent with the latter concept, both *let-92* RNAi and *sur-6* RNAi impaired SPD-2 and PLK-1 departure from PCM during anaphase (Figure S4D-G) (Magescas et al., 2019).

**Figure 6.**
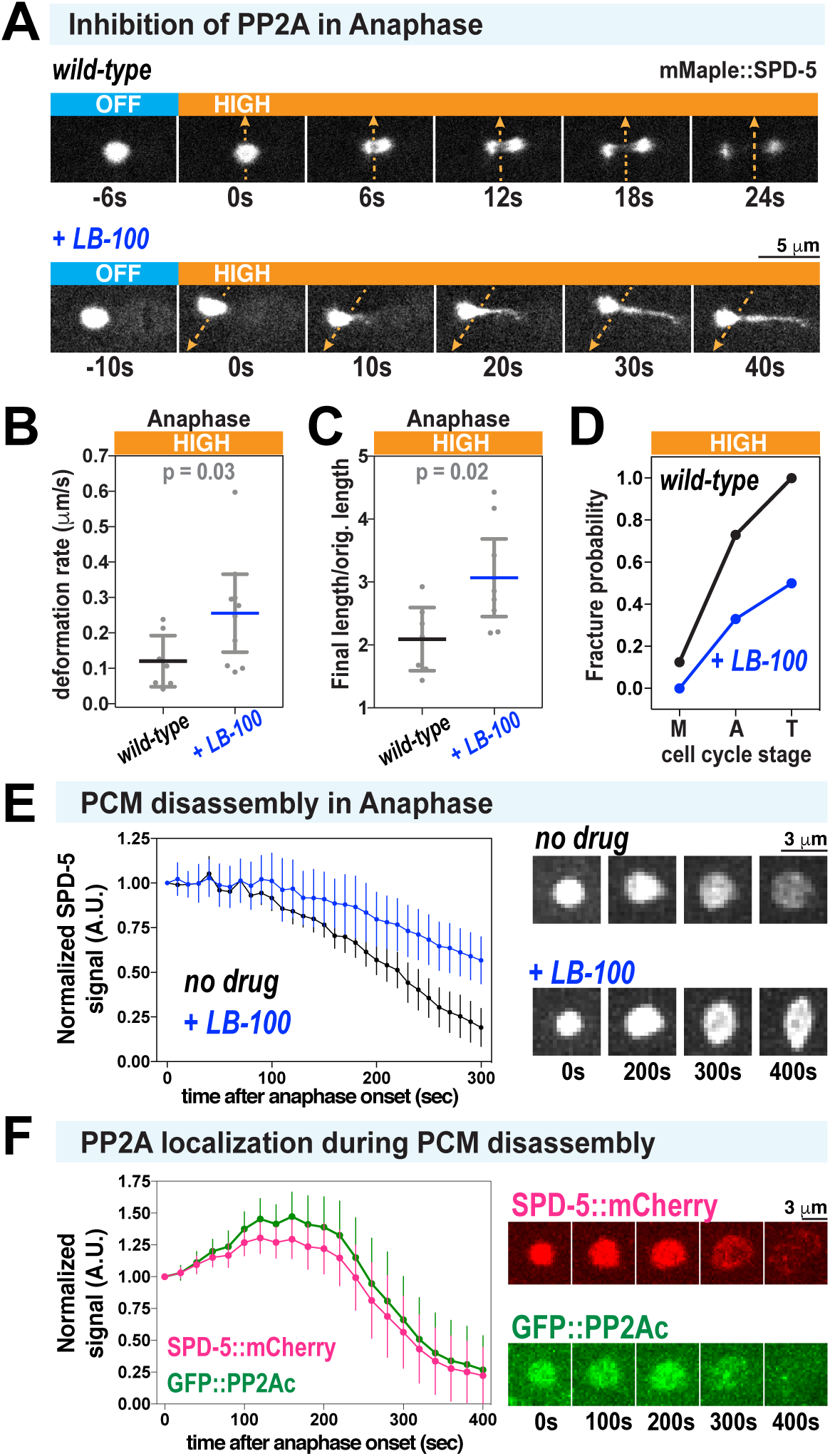
PCM becomes fracture-resistant and ductile in anaphase after inhibition of PP2A phosphatase. A. PCM was subjected to high-flow FLUCS during anaphase in wild-type embryos or permeabilized embryos treated with 10 μM LB-100 (inhibitor of PP2A phosphatase). B. PCM deformation rates in anaphase using high flow in wild-type and LB-100-treated embryos. Wild-type data are from experiments in Figure 1. Individual data points are plotted with bars representing mean +/− 95% CI; n = 7 (wild-type) and 10 (LB-100-treated) centrosomes. P-values were calculated using a Mann-Whitney test. C. Ratio of final PCM length to original length in experiments from (B). Original PCM length was measured before flow began and final PCM length was measured once flow was turned off. P-values were calculated using a Mann-Whitney test. D. PCM fracture probabilities for high-flow FLUCS in metaphase (M), anaphase (A), and telophase (T). Wild-type data are from experiments in Figure 1; n= 8,11,11 (wild-type) and 7,9,6 (LB-100-treated) centrosomes. E. Permeabilized embryos were treated with no drug or 10 μM LB-100 in metaphase, then imaged until 300s after anaphase onset. Data are plotted as normalized lines representing mean +/− 95% CI; n = 24 (no drug) and n = 21 (LB-100-treated) centrosomes. F. Dual-color imaging of embryos expressing GFP-tagged LET-92, the PP2A catalytic subunit in *C. elegans* (GFP::PP2Ac), and SPD-5::mCherry. Data are plotted as normalized lines representing mean +/− 95% CI; n = 10 centrosomes.

To determine when and where PP2A might dephosphorylate PCM proteins, we visualized embryos expressing GFP::LET-92, the PP2A catalytic subunit in *C. elegans* (Schlaitz et al., 2007). GFP::LET-92 localized to the PCM and persisted there until SPD-5 scaffold disassembly, approximately 100s after PLK-1 had departed from the PCM (Figure 6F), consistent with previous observations (Magescas et al., 2019). Our results suggest that, during anaphase, Polo Kinase activity at the PCM ceases and PP2A removes Polo-delivered phosphates and contributes to SPD-2 departure. This shift in the balance of phosphorylation and dephosphorylation changes the mechanical properties of the PCM, making it more brittle and susceptible to fracture and dissolution.

## DISCUSSION

Mitotic spindle assembly and positioning require that centrosomes bear tensile microtubule-dependent forces without structural failure. As mitosis ends, however, these same forces are sufficient to deform and fracture centrosomes, facilitating their disassembly. Disassembly is essential to release centrioles and avoid accumulation of old centrosomes over successive rounds of cell division. Here, we combined flow-driven mechanical perturbations *in vivo* with biochemical reconstitution *in vitro* to determine the molecular mechanisms regulating deformation resistance and fracture resistance of PCM, the outer and most massive layer of a centrosome.

### PCM mechanical properties, function, and renewal can be achieved through transient reinforcement of the PCM scaffold

Using *C. elegans* embryos as a model system, we found that PCM deformation resistance, fracture resistance, and composition are tuned in a cell-cycle-dependent manner (Figure 7A,B). During metaphase, PCM resists both microtubule-mediated forces and induced flow perturbations without deforming or fracturing. In this state, PCM is structured as a reinforced composite, comprising a non-dynamic scaffold of SPD-5 molecules filled with a dynamic phase of regulatory molecules like SPD-2 and PLK-1, which frequently bind and unbind the scaffold (Figure 7B). During anaphase, PCM loses PLK-1 and SPD-2 and then becomes susceptible to deformation and fracture. During telophase, PCM is at its weakest and is easily fractured and dispersed by microtubule-mediated forces, a hallmark step in the PCM disassembly process.

**Figure 7.**
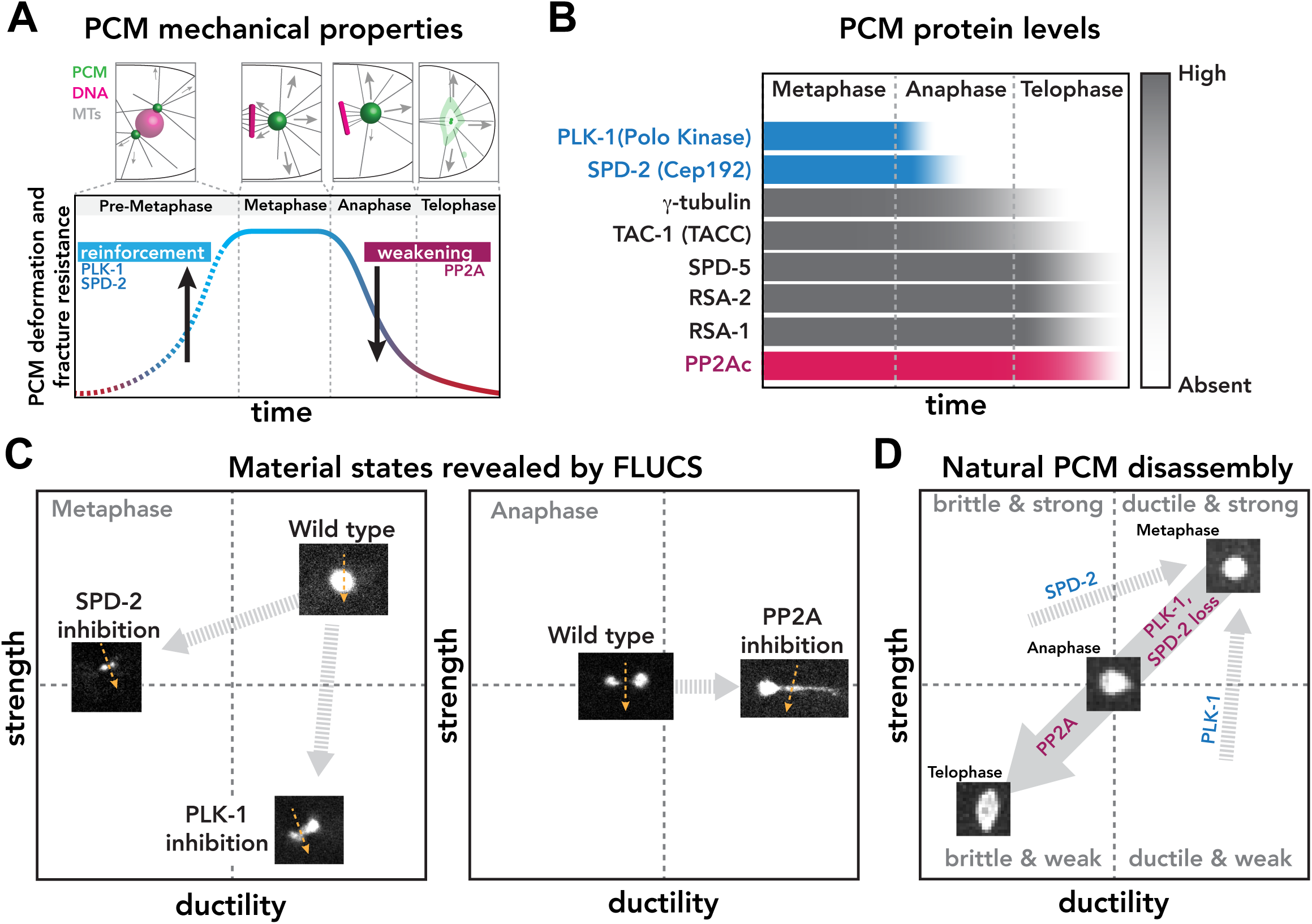
The balance of PLK-1, SPD-2, and PP2A activities tune PCM strength and ductility. A. PCM resistance to microtubule-mediated forces peaks in metaphase during spindle assembly, then declines in anaphase and telophase, corresponding to PCM disassembly. B. PCM-localized levels of 8 different proteins during mitotic progression. In anaphase, PLK-1 levels decline first, followed by SPD-2. The catalytic subunit of PP2A phosphatase (PP2Ac), as well as the main scaffold protein SPD-5, remain at the PCM until late telophase. C. In metaphase, FLUCS cannot fracture or deform wild-type PCM. However, FLUCS can fracture PCM in *spd-2* mutant embryos (i.e., PCM is less ductile) and stretch and deform PCM in PLK-1 inhibited embryos (i.e., PCM is less strong but still ductile). In anaphase, FLUCS easily deforms and fractures wild-type PCM, while it deforms and stretches PCM in PP2A-inhibited embryos (i.e., PCM is more ductile). D. The combination of PP2A phosphatase activity and the departure of PLK-1 and SPD-2 transitions PCM from a strong, ductile state in metaphase to a weak, brittle state in telophase. This transition enables PCM disassembly and dispersal through microtubule-mediated pulling forces.

Our implementation of flow perturbations *in vivo* using FLUCS reveals how PLK-1, SPD-2, and PP2A contribute to the dynamic mechanical properties of the PCM. We interpret deformation resistance as an indicator of strength and fracture resistance and strain as indicators of ductility. Wild-type metaphase PCM is highly resistant to flow perturbations, but underlying features appear in different mutant states (Figure 7C). When PLK-1 is inhibited, the PCM scaffold is easily deformed by FLUCS and stretches without fracturing. Thus, PLK-1 normally maintains PCM strength. On the other hand, when SPD-2 is inhibited, the PCM scaffold is easily fractured and dissolved by FLUCS but does not stretch. Thus, SPD-2 normally maintains PCM ductility and strength. Elimination of either PLK-1 or SPD-2 causes premature PCM disassembly, suggesting that the combination of strength and ductility is necessary for PCM function and maintenance during spindle assembly in metaphase. In anaphase, wild-type PCM is more easily deformed and fractured by FLUCS. Yet, when PP2A is inhibited, PCM is difficult to fracture by FLUCS and instead stretches up to 4 times its original length, revealing that the high ductility of the PCM, which was established prior to metaphase, is preserved. Thus, PP2A normally functions to drive PCM disassembly by reducing PCM ductility.

We propose that that the balance of PLK-1, SPD-2, and PP2A activities determine the mechanical properties and assembly/disassembly state of the PCM (Figure 7D). In metaphase, PLK-1 phosphorylation of SPD-5 and direct binding of SPD-2 reinforce the SPD-5 scaffold, conferring strength and ductility. During anaphase, PLK-1 and SPD-2 depart from the PCM, while PP2A phosphatase remains and removes PLK-1-derived phosphates. As a result, PCM becomes progressively brittle and weak, allowing microtubule-dependent forces to deform and fracture it in telophase. Since PLK-1 and SPD-2 stabilize the SPD-5 scaffold, but are themselves dynamic, we call this mode of PCM regulation “transient reinforcement”.

Transient reinforcement of the PCM scaffold, in theory, could enable cell-cycle regulated PCM assembly, function, and renewal. In preparation for mitosis, PCM must rapidly assemble and provide a solid foundation for nucleating and anchoring microtubules. If PCM assembly fails, then mitotic spindle assembly and chromosome segregation is severely impaired (Doxsey et al., 1994; Hamill et al., 2002; Sunkel and Glover, 1988). PLK-1 and SPD-2 thus play dual roles in PCM functionality: 1) catalyzing assembly of the PCM scaffold and 2) strengthening it to withstand microtubule-dependent pulling forces (shown here). While PCM is stable during spindle assembly, PCM disassembles in telophase only to be rebuilt in the next cell cycle. PCM disassembly is essential for entry into various post-mitotic states, including the formation of acentriolar oocytes and heart tissue (Pimenta-Marques et al., 2016; Zebrowski et al., 2015). How might transient reinforcement enable PCM disassembly and renewal? Based on *in vivo* and *in vitro* FRAP data, PLK-1 and SPD-2 are mobile within the PCM, suggesting that they frequently bind and unbind the SPD-5 scaffold (Laos et al., 2015; Woodruff et al., 2017). Either decreasing their association or increasing their dissociation rates with the SPD-5 scaffold would reduce SPD-2 and PLK-1 levels at the PCM. One potential mechanism is through localized ubiquitination and degradation, which controls Polo Kinase levels at centrosomes in human tissue culture cells during anaphase (Lindon and Pines, 2004). It is currently unknown how PCM levels of SPD-2 or its homolog Cep192 are tuned. The completeness and speed of SPD-2 and PLK-1 removal during anaphase is also suggestive of feedback. Thus, the system could be set up such that minor changes to SPD-2 and PLK-1 affinity elicit large, switch-like changes in PCM structure and mechanical properties. Further experiments are needed to define how the other numerous PCM proteins and centriole tethers contribute to PCM mechanical properties. One possible control point is linkage of the SPD-5 scaffold to the centriole via PCMD-1; inactivation of PCMD-1 or laser ablation of centrioles leads to aberrant PCM deformation, presumably because PCM can no longer fully resist microtubule-pulling forces (Cabral et al., 2019; Erpf et al., 2019).

### PCM mechanical properties in other eukaryotes

The mechanical properties of PCM in other species have yet to be determined. We speculate that our transient reinforcement model for tuning PCM load bearing capacity may be conserved for three reasons. First, diverse eukaryotic species—nematodes, frogs, flies, and human cells—use both Polo Kinase and SPD-2/Cep192 to enhance assembly of the PCM scaffold (Conduit et al., 2014b; Decker et al., 2011; Haren et al., 2009; Joukov et al., 2014; Kemp et al., 2004; Pelletier et al., 2004; Woodruff et al., 2015). Second, PP2A is highly conserved in eukaryotes and is required for mitotic exit in these species (Wlodarchak and Xing, 2016). Third, in *Drosophila* embryos, PCM undergoes cell-cycle-regulated deformation and fracture, termed “flaring”, which appears similar to disassembling PCM in *C. elegans* (Megraw et al., 2002). PCM flares are visible during interphase, cease during metaphase and anaphase, and then escalate during telophase. Flares also require dynamic microtubules. Thus, *Drosophila* PCM flaring may be due to decreasing PCM strength and ductility during telophase, such that PCM can no longer resist microtubule-mediated forces.

### Parallels between PCM and common soft materials in engineering

The mechanical properties and structure of mature PCM are analogous to common composite materials such as flexible plastics and hydrogels. Most modern plastics comprise cross-linked polymer chains embedded with plasticizers, chemicals that make the plastic more flexible and ductile. Over time, these plasticizers exit by diffusion, making the remaining plastic brittle and weak, which is a form of material aging. The rubber sole on a shoe will crack with age; flexibility of the old sole can be restored by impregnating the rubber with a plasticizer such as silicone. For PCM, the SPD-5 scaffold is most similar to the polymer chains, whereas PLK-1 and SPD-2 could act as plasticizers. Similar to aging rubber losing its plasticizers, our results show that the PCM scaffold becomes brittle and weak during anaphase, coincident with both PLK-1 and SPD-2 leaving the PCM.

PCM is also similar to physical composites like polyampholyte hydrogels, which exhibit a unique combination of high tensile strength and flexibility. Polyampholyte gels comprise polymers cross-linked with a combination of high- and low-affinity non-covalent bonds. Upon stress, the low-affinity bonds break and dissipate energy, while the high-affinity bonds maintain the overall supramolecular structure. The low affinity bonds then quickly reform, resulting in self-healing that prevents structural fatigue from repeated stresses (Sun et al., 2013). These bonds also make the material more ductile, such that it will undergo plastic deformation instead of fracturing. For PCM, it is possible that PLK-1, SPD-2, and other PCM-resident proteins dissipate stress by unbinding from the PCM scaffold, then re-binding to achieve self-healing. Eliminating these weak interactions would make the PCM weaker and more brittle. This would naturally occur in anaphase as PLK-1 and SPD-2 depart from PCM. This concept could also explain why FLUCS induces PCM fracture and deformation in metaphase when we acutely inhibit PLK-1 and SPD-2. Alternatively, in PP2A-inhibited embryos, PCM is ductile due to the preservation of low affinity, self-healing bonds. Although stronger and more ductile than normal, this mutant PCM still weakens and disassembles in telophase, suggesting that another yet unknown process disrupts the strong interactions between SPD-5 molecules.

## Conclusion

This work establishes that PCM, the most substantial layer of the centrosome, transitions from a strong, ductile state in metaphase to a weak, brittle state in telophase. This transition is driven by PP2A phosphatase and inactivation of Polo Kinase and SPD-2/Cep-192, which are essential for centrosome assembly and reinforce the PCM scaffold during metaphase. This mode of mechanical regulation, which we term “transient reinforcement”, is a functional form of material aging that allows PCM to resist microtubule-mediated tensile stresses during spindle assembly and then to be fractured and disassembled by similar forces during mitotic exit. Implicitly, our work demonstrates how flow perturbations can reveal functional mechanical states of membrane-less organelles in vivo.

## ACKNOWLEDGEMENTS

We would like to thank Karen Oegema, Arshad Desai, Anthony Hyman, and Bruce Bowerman for providing strains; Andrea Zinke, Anne Schwager, and Susanne Ernst for help with worm maintenance; Carsten Hoege and David Drechsel for providing the CRISPR reagents; Stephan Daetwyler and Robert Ernst for help with image analysis and script writing; the Live Cell Imaging Facility at UT Southwestern and Light Microscopy Facility at the MPI-CBG for help with microscopy; the Protein Expression Facility at MPI-CBG for help with protein expression.

## AUTHOR CONTRIBUTIONS

M.M., A.W.F., and M.K. built the FLUCS microscope. M.M. and V.M.T. performed and analyzed the FLUCS experiments on *C. elegans* embryos. M.U.R. performed the *spd-2* temperature-sensitive experiments. B.F-G. and S.J.E. performed two-color imaging of PCM proteins during anaphase and wrote the image analysis scripts. A.B. performed imaging of *sur-6(RNAi)* mutants. J.B.W. purified proteins and performed the *in vitro* assays. V.M.T. performed all other in vivo experiments and analysis. J.B.W. and M.K. analyzed data and wrote the manuscript.

## SUPPLEMENTAL FIGURE LEGENDS

**Figure S1.**
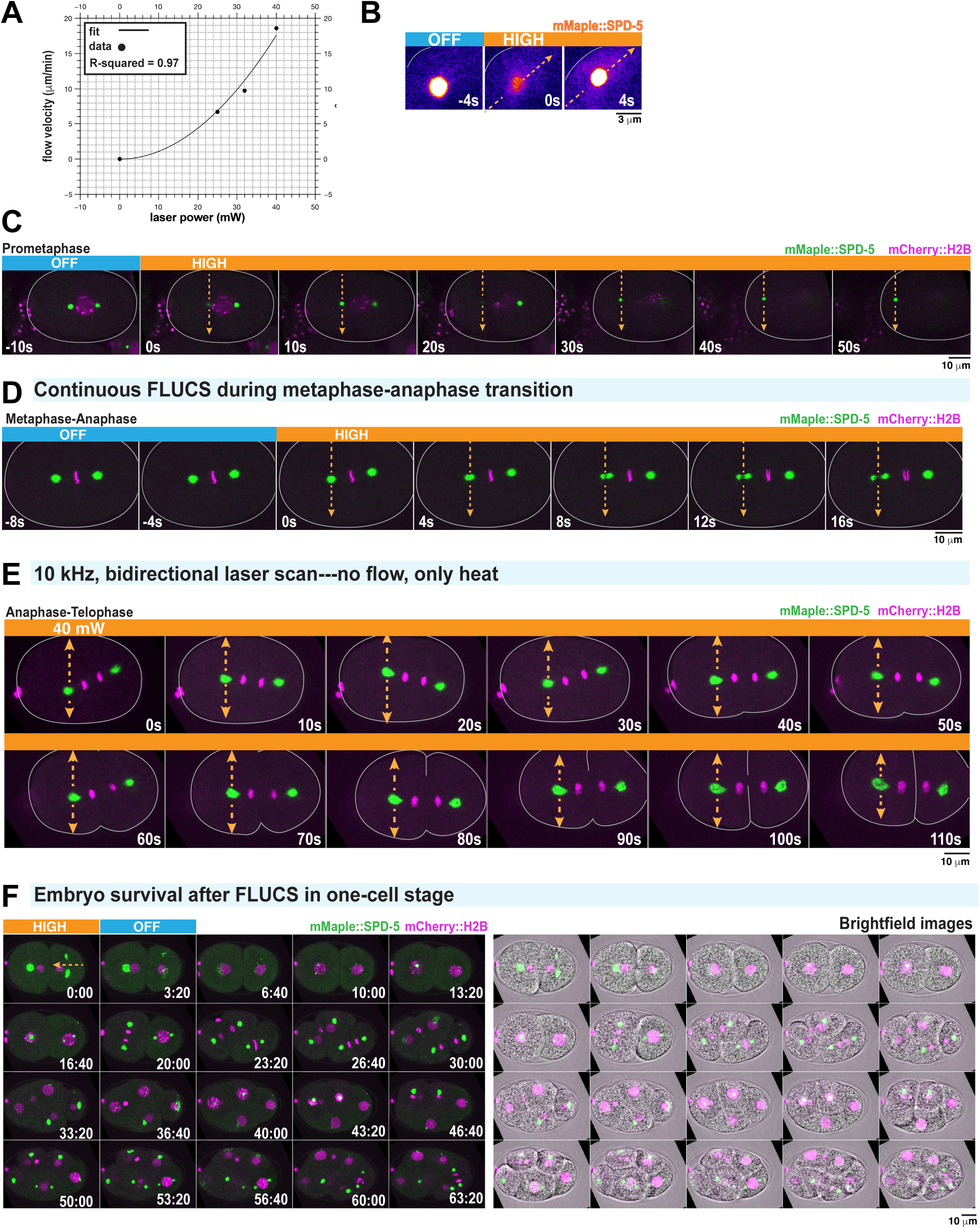
FLUCS control experiments. A. Power law scaling of cytoplasmic flow with increasing FLUCS laser power. Individual data points represent mean +/− 95% CI; n = 5 embryos per laser condition. Flow velocities were fit with a second-order polynomial. B. Application of high-flow FLUCS in an anaphase 1-cell embryo. Images are pseudo-colored to highlight the subtle increase in cytoplasmic mMaple::SPD-5 fluorescence after flow begins. Note: the centrosome goes out of focus in the first frame when FLUCS begins. C. Application of high-flow FLUCS in a prometaphase 1-cell embryo. Single plane images are shown. Flow causes the centrosome to leave the plane of focus at t=0s and t=20s. Flow then displaces the centrosome toward the cortex. D. High-flow FLUCS was applied in metaphase, continuing into anaphase (indicated by chromosome segregation at t=12s). E. Example images from the experiment in Figure 1H. Bidirectional scanning of a 40 mW laser (1455 nm) at 10 kHz creates local temperature gradients without generating flow. F. Time-lapse fluorescence and brightfield images of an embryo after cessation of FLUCS. Application of high-flow FLUCS does not affect embryonic development.

**Figure S2.**
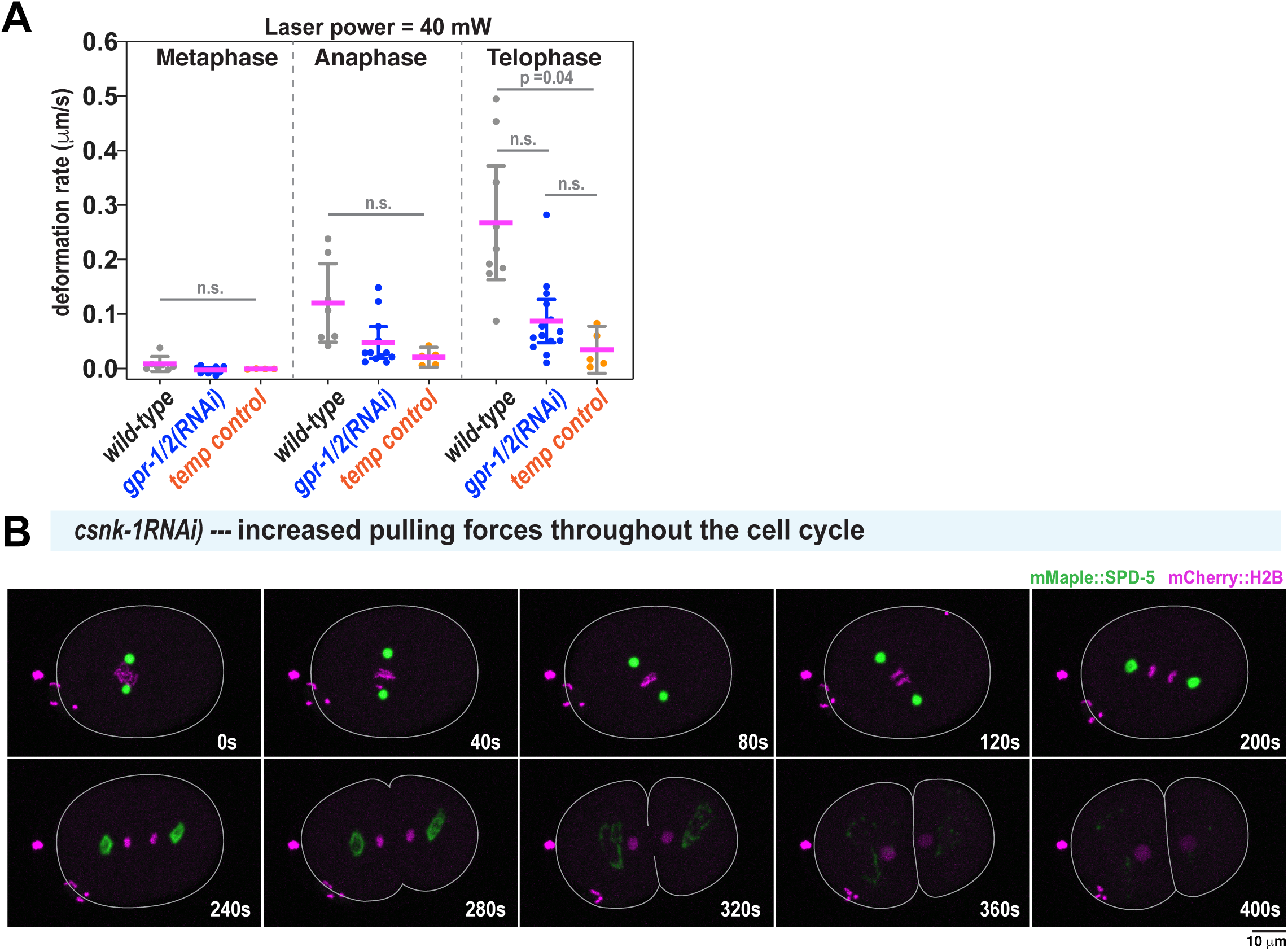
Contributions of microtubule pulling forces to PCM deformation. A. PCM deformation rates in metaphase (M), anaphase (A), and telophase (T) using high flow in wild-type and *gpr-1/2(RNAi)* embryos or 40 mW bidirectional laser scanning (temperature control; no flow). Data are from experiments in Figures 1 and 2. Individual data points are plotted with mean +/− 95% CI; n = 7,7,9 (wild-type; metaphase, anaphase, telophase), n = 10,12,13 (*gpr-1/2(RNAi)* and n = 4,5,5 (temperature control). P values were calculated using Brown-Forsythe and Welch ANOVA tests followed by Dunnett’s T3 multiple comparisons tests. B. Time-lapse images of centrosomes in a *csnk-1(RNAi)* embryo, where microtubule-mediated pulling forces at the cortex are ∼1.5-fold elevated compared to wild type (Panbianco et al., 2008). PCM deformation does not occur prematurely in metaphase.

**Figure S3.**
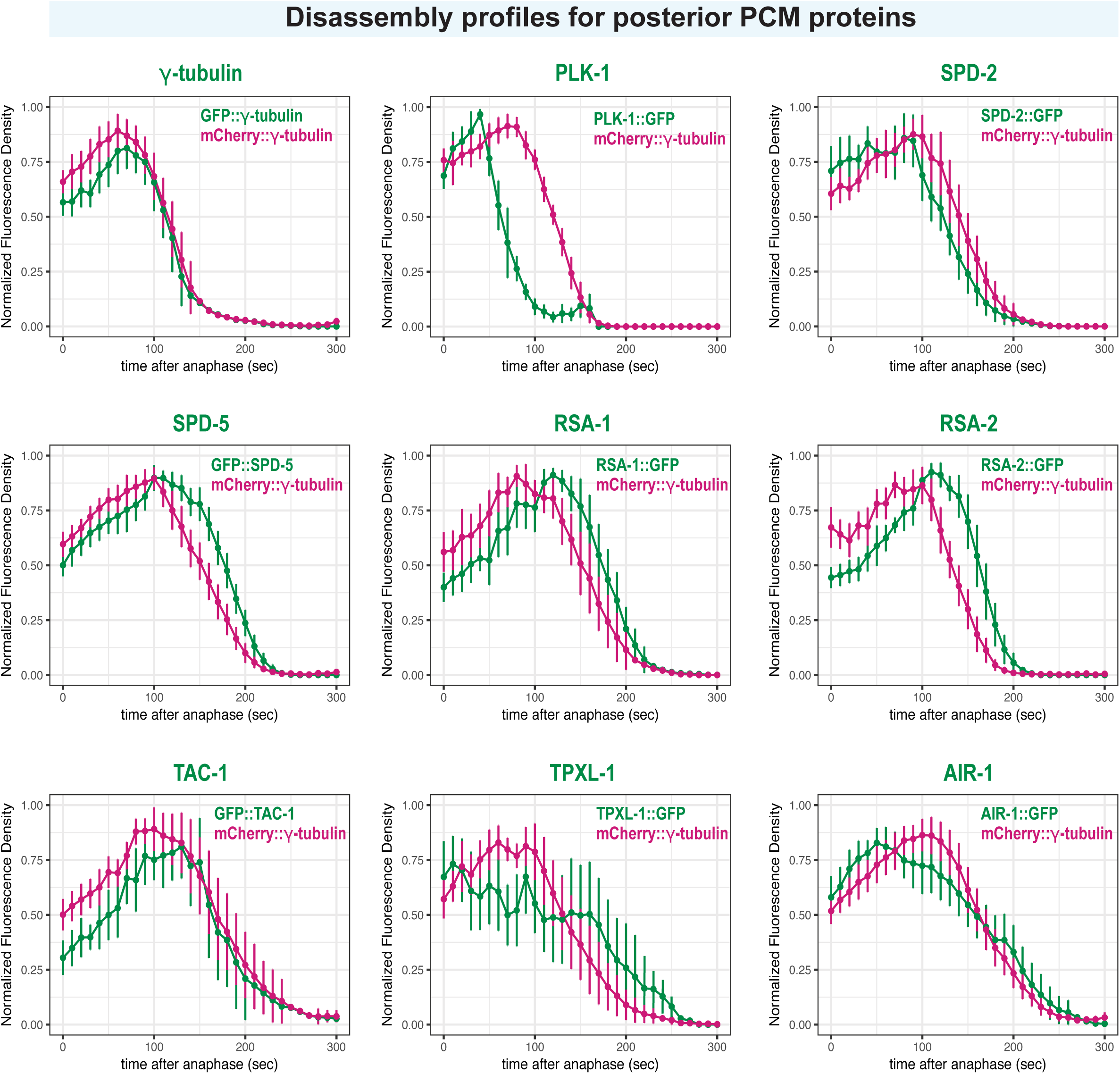
Localization profiles of PCM proteins in the posterior embryo during PCM disassembly. Quantification of PCM localization in the posterior side of 1-cell embryos in Figure 3. For each strain, the plots represent the normalized integrated fluorescence density of PCM-localized mCherry-tagged *γ*-tubulin compared to the GFP-tagged protein from anaphase onward. Anaphase was indicated by spindle rocking. Data are normalized to maxima for each individual curve, then averaged; mean +/− 95% C.I. n = 13 (*γ*-tubulin), 9 (PLK-1), 10 (SPD-2), 12 (SPD-5), 14, (RSA-1), 12 (RSA-2), 7 (TAC-1), 10 (TPXL-1), 13 (AIR-1)).

**Figure S4.**
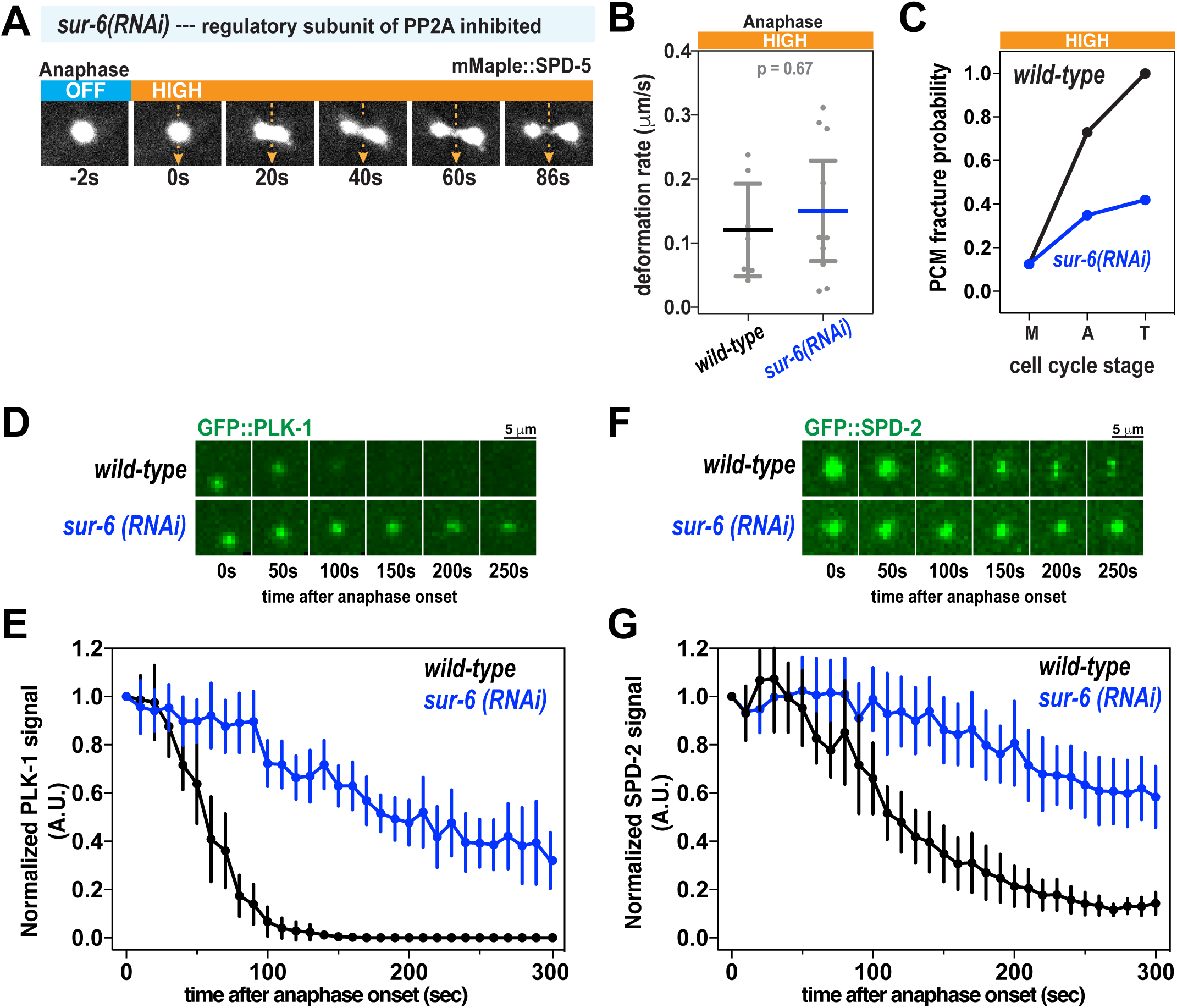
FLUCS and localization experiments in *sur-6(RNAi)* embryos. A. High-flow FLUCS was applied to a centrosome in an embryo depleted of SUR-6, a PP2A regulatory subunit involved in PCM disassembly. B. PCM deformation rates in anaphase during high-flow FLUCS in wild-type and *sur-6(RNAi)* embryos. Wild-type data are from experiments in Figure 1. Individual data points are plotted with bars representing mean +/− 95% CI; n = 7 (wild-type) and 10 (*sur-6(RNAi)*) centrosomes. C. PCM fracture probabilities in metaphase (M), anaphase (A), and telophase (T) during high-flow FLUCS experiments. N= 8-11 (wild-type) and 8-17 (*sur-6(RNAi)*) centrosomes. D. Images of PCM-localized GFP::PLK-1 in wild-type and *sur-6(RNAi)* embryos. E. Normalized integrated fluorescence intensity of PCM-localized GFP::SPD-2 during anaphase. Data are plotted as mean +/− 95% CI; n= 22 (wild-type) and 32 (*sur-6(RNAi)*) centrosomes. F. Images of PCM-localized GFP::SPD-2 in wild-type and *sur-6(RNAi)* embryos. G. Normalized integrated fluorescence intensity of PCM-localized GFP::SPD-2 during anaphase. Data are plotted as mean +/− 95% CI; n= 20 (wild-type) and 24 (*sur-6(RNAi)*) centrosomes.

## SUPPLEMENTAL MOVIES

**Movie S1. FLUCS flow control using 25 mW, 32 mW, and 40 mW laser scans at 1.5 kHz.** Flows were generated in *C. elegans* 1-cell embryos using three different 1455 nm laser powers (25 mW, 32 mW, and 40 mW).

**Movie S2. High-flow FLUCS targeting the centrosome in a prometaphase 1-cell embryo.** Prometaphase *C. elegans* embryos expressing mCherry::histoneH2B (magenta) and GFP::SPD-5 (green) were subjected to high-flow FLUCS (40 mW). Images are of a single confocal plane.

**Movie S3. High-flow FLUCS targeting the centrosome in a metaphase 1-cell embryo.** Metaphase *C. elegans* embryos expressing mCherry::histoneH2B (magenta) and GFP::SPD-5 (green) were subjected to high-flow FLUCS (40 mW). Images are of a single confocal plane.

**Movie S4. High-flow FLUCS targeting the centrosome in an anaphase 1-cell embryo.** Anaphase *C. elegans* embryos expressing mCherry::histoneH2B (magenta) and GFP::SPD-5 (green) were subjected to high-flow FLUCS (40 mW). Images are of a single confocal plane.

**Movie S5. High-flow FLUCS targeting the centrosome in a telophase 1-cell embryo.** Telophase *C. elegans* embryos expressing mCherry::histoneH2B (magenta) and GFP::SPD-5 (green) were subjected to high-flow FLUCS (40 mW). Images are of a single confocal plane.

**Movie S6. High-flow FLUCS targeting the centrosome during the metaphase to anaphase transition in a 1-cell embryo.** *C. elegans* embryos expressing mCherry::histoneH2B (magenta) and GFP::SPD-5 (green) were subjected to high-flow FLUCS (40 mW) during the metaphase-anaphase transition. Images are of a single confocal plane.

**TABLE S1.** C. elegans strains used in this study.

| Strain name | genotype | Creation method | Origin |
| --- | --- | --- | --- |
| DAM858 | vie11[pAD676; gfp::tac-1]II | CRISPR | Alexander Dammermann |
| EU584 | spd-2(or188ts) I | mutagenesis | Bruce Bowerman |
| JWW1 | utsw2[mMaple::spd-5] I | CRISPR | This study |
| JWW13 | spd-2(or188ts) I; unc-119(ed9) III; ltSi202[pVV103/pOD1021; Pspd-2::GFP::SPD-5 RNAiresistant;cb-unc-119(+)]II | Cross of EU584 and OD847 | This study |
| JWW35 | ltSi202[pVV103/ pOD1021; Pspd-2::GFP::SPD-5 RNAiresistant;cb-unc-119(+)]II ; unc-119(ed3)III; ddls44[WRM0614cB02 GLCherry::tbg-1;Cbr-unc-119(+)] | Cross of OD847 and TH169 | This study |
| JWW64 | utsw2[mMaple::spd-5] I; ltIs37 [(pAA64) pie-1p::mCherry::his-58 + unc-119(+)] IV. | Cross of JWW1 and OD95 | This study |
| JWW65 | lt17[plk-1::gfp+loxP]III ; ltIs37 [(pAA64) pie-1p::mCherry::his-58 + unc-119(+)] IV; unc-119(ed3) III | Cross of OD2425 and OD95 | This study |
| JWW66 | ltSi203[pVV60; Pspd-2::GFP::SPD-2 reencoded; cb-unc-119(+)]II; ltIs37 [(pAA64) pie-1p::mCherry::his-58 + unc-119(+)] IV; unc-119(ed3) III | Cross of OD824 and OD95 | This study |
| JWW67 | unc-119(ed9) III; utsw1[pJWB56; Pspd-2::GFP::SPD-5(530E, 627E, 653E, 658E) re-encoded; cb-unc-119(+)]II | MosSCI, into EG6699 | This study |
| JWW69 | unc-119(ed9) III; ltSi202[pVV103/ pOD1021; Pspd-2::GFP::SPD-5 RNAiresistant;cb-unc-119(+)]II; ltIs37 [(pAA64) pie-1p::mCherry::his-58 + unc-119(+)] IV. | Cross of OD847 and OD95 | This study |
| JWW70 | unc-119(ed9) III; utsw1[pJWB56; Pspd-2::GFP::SPD-5(530E, 627E, 653E, 658E) re-encoded; cb-unc-119(+)]II; ltIs37 [(pAA64) pie-1p::mCherry::his-58 + unc-119(+)] IV. | Cross of JWW1 and OD95 | This study |
| JWW71 | lt17[plk-1::gfp+loxP]III; unc-119(ed3)III; ddIs44[WRM0614cB02 GLCherry::tbg-1;Cbr-unc-119(+)] | Cross of OD2425 and TH169 | This study |
| JWW72 | vie11[pAD676; gfp::tac-1]II ; unc-119(ed3)III; ddIs44[WRM0614cB02 GLCherry::tbg-1;Cbr-unc-119(+)] | Cross of DAM858 and TH169 | This study |
| JWW89 | spd-2(or188ts) I; ltSi202[pVV103/ pOD1021; Pspd-2::GFP::SPD-5 RNAiresistant;cb-unc-119(+)]II; ltIs37 [(pAA64) pie-1p::mCherry::his-58 + unc-119(+)] IV. | Cross of JWW13 and OD95 | This study |
| OD2425 | lt17[plk-1::gfp+loxP]III | CRISPR | Karen Oegema |
| OD823 | ltSi203[pVV60; Pspd-2::GFP::SPD-2 reencoded; cb-unc-119(+)]II; unc-119(ed3) III | MosSCI, into EG6699 | Karen Oegema |
| OD847 | unc-119(ed9) III; ltSi202[pVV103/ pOD1021; Pspd-2::GFP::SPD-5 RNAiresistant;cb-unc-119(+)]II | MosSCI, into EG6699 | (Woodruff et al., 2015) |
| OD95 | unc-119(ed3) III; ltIs37 [(pAA64) pie-1p::mCherry::his-58 + unc-119(+)] IV; ltIs38 [pie-1p::GFP::PH(PLC1delta1) + unc-119(+)] | Microparticle bombardment | CGC |
| TH169 | unc-119(ed3)III; ddIs44[WRM0614cB02 GLCherry::tbg-1;Cbr-unc-119(+)] | Microparticle bombardment | Anthony Hyman |
| TH447 | unc-119(ed9) III; ddIs243[pie-1p::LAP::LET-92; unc-119(+)] ; ddIs247[pie-1p::SPD-5(synthetic introns, CAI 0.65)::mCherry; unc-119(+)] | Microparticle bombardment | Anthony Hyman |
| TH530 | rsa-1::LAP; unc-119(ed3)III; ddIs44[WRM0614cB02 GLCherry::tbg-1;Cbr-unc-119(+)] | Microparticle bombardment | Anthony Hyman |
| TH531 | rsa-2::LAP; unc-119(ed3)III; ddIs44[WRM0614cB02 GLCherry::tbg-1;Cbr-unc-119(+)] | Microparticle bombardment | Anthony Hyman |
| TH539 | spd-2::GFP; unc-119(ed3)III; ddIs44[WRM0614cB02 GLCherry::tbg-1;Cbr-unc-119(+)] | Microparticle bombardment | Anthony Hyman |
| TH571 | unc-119(ed3)III; ddIs12[pie-1p::tpxl-1::GFP;unc-119(+)] ; ddIs44[WRM0614cB02 GLCherry::tbg-1;Cbr-unc-119(+)] | Microparticle bombardment | Anthony Hyman |
| TH630 | ddIs44[WRM0614cB02 GLCherry::tbg-1;Cbr-unc-119(+)] ; ddIs62[pie-1p::AIR-1(synthetic introns, CAI 1.0)::GFP; unc-119(+)] ; unc-119(ed3)III | Microparticle bombardment | Anthony Hyman |
| EG6699 | ttTi5605 II; unc-119(ed3) III; oxEx1578. |  | CGC |

**TABLE S2.** Protein expression plasmids used in this study.

| Plasmid name | Gene | N-term tag | C-term tag | Origin |
| --- | --- | --- | --- | --- |
| JWV11 | *plk-1(T194D)*<br>constitutively active |  | PreScission-6xHis | (Woodruff et al., 2015) |
| JWV12 | *plk-1(K67M)*<br>kinase dead |  | PreScission-6xHis | (Woodruff et al., 2015) |
| JWV2 | *spd-5(wt)* | MBP-PreScission | PreScission-6xHis | (Woodruff et al., 2015) |
| JWV3 | *spd-5(wt)* | MBP-PreScission | tagRFP-PreScission-6xHis | (Woodruff et al., 2015) |
| JWV6 | *spd-2(wt)* | MBP-TEV | TEV-6xHis | (Woodruff et al., 2015) |

**TABLE S3.** One-way ANOVA and post-hoc tests of anterior PCM disassembly profiles from Figure 3F.

| Holm-Sidak's multiple comparisons test | Mean Diff. | Significant? | Summary | Adjusted P Value |
| --- | --- | --- | --- | --- |
| AIR-1 vs. TPXL-1 | -25.55 | No | ns | 0.2725 |
| AIR-1 vs. RSA-1 | -8.310 | No | ns | 0.9615 |
| AIR-1 vs. RSA-2 | -14.38 | No | ns | 0.8174 |
| AIR-1 vs. SPD-2 | 38.26 | Yes | ** | 0.0079 |
| AIR-1 vs. SPD-5 | -24.73 | No | ns | 0.2183 |
| AIR-1 vs. TBG-1 | 8.728 | No | ns | 0.9598 |
| AIR-1 vs. TAC-1 | 0.4412 | No | ns | 0.9963 |
| AIR-1 vs. PLK-1 | 92.29 | Yes | **** | <0.0001 |
| TPXL-1 vs. RSA-1 | 17.24 | No | ns | 0.8060 |
| TPXL-1 vs. RSA-2 | 11.16 | No | ns | 0.9598 |
| TPXL-1 vs. SPD-2 | 63.81 | Yes | **** | <0.0001 |
| TPXL-1 vs. SPD-5 | 0.8171 | No | ns | 0.9963 |
| TPXL-1 vs. TBG-1 | 34.28 | Yes | * | 0.0356 |
| TPXL-1 vs. TAC-1 | 25.99 | No | ns | 0.4307 |
| TPXL-1 vs. PLK-1 | 117.8 | Yes | **** | <0.0001 |
| RSA-1 vs. RSA-2 | -6.074 | No | ns | 0.9615 |
| RSA-1 vs. SPD-2 | 46.57 | Yes | ** | 0.0012 |
| RSA-1 vs. SPD-5 | -16.42 | No | ns | 0.8060 |
| RSA-1 vs. TBG-1 | 17.04 | No | ns | 0.7693 |
| RSA-1 vs. TAC-1 | 8.751 | No | ns | 0.9615 |
| RSA-1 vs. PLK-1 | 100.6 | Yes | **** | <0.0001 |
| RSA-2 vs. SPD-2 | 52.64 | Yes | **** | <0.0001 |
| RSA-2 vs. SPD-5 | -10.35 | No | ns | 0.9598 |
| RSA-2 vs. TBG-1 | 23.11 | No | ns | 0.2887 |
| RSA-2 vs. TAC-1 | 14.83 | No | ns | 0.8971 |
| RSA-2 vs. PLK-1 | 106.7 | Yes | **** | <0.0001 |
| SPD-2 vs. SPD-5 | -62.99 | Yes | **** | <0.0001 |
| SPD-2 vs. TBG-1 | -29.53 | No | ns | 0.0940 |
| SPD-2 vs. TAC-1 | -37.82 | Yes | * | 0.0454 |
| SPD-2 vs. PLK-1 | 54.03 | Yes | *** | 0.0002 |
| SPD-5 vs. TBG-1 | 33.46 | Yes | * | 0.0207 |
| SPD-5 vs. TAC-1 | 25.17 | No | ns | 0.4085 |
| SPD-5 vs. PLK-1 | 117.0 | Yes | **** | <0.0001 |
| TBG-1 vs. TAC-1 | -8.286 | No | ns | 0.9615 |
| TBG-1 vs. PLK-1 | 83.56 | Yes | **** | <0.0001 |
| TAC-1 vs. PLK-1 | 91.85 | Yes | **** | <0.0001 |

**TABLE S4.** One-way ANOVA and post-hoc tests of posterior PCM disassembly profiles from Figure 3G.

| Holm-Sidak's multiple comparisons test | Mean Diff. | Significant? | Summary | Adjusted P Value |
| --- | --- | --- | --- | --- |
| AIR-1 vs. TPXL-1 | -9.235 | No | ns | 0.9002 |
| AIR-1 vs. RSA-1 | -36.70 | Yes | *** | 0.0003 |
| AIR-1 vs. RSA-2 | -46.04 | Yes | **** | <0.0001 |
| AIR-1 vs. SPD-2 | 5.014 | No | ns | 0.9652 |
| AIR-1 vs. SPD-5 | -42.30 | Yes | **** | <0.0001 |
| AIR-1 vs. TBG-1 | -13.68 | No | ns | 0.5679 |
| AIR-1 vs. TAC-1 | -34.04 | Yes | ** | 0.0041 |
| AIR-1 vs. PLK-1 | 40.38 | Yes | **** | <0.0001 |
| TPXL-1 vs. RSA-1 | -27.46 | Yes | * | 0.0334 |
| TPXL-1 vs. RSA-2 | -36.81 | Yes | *** | 0.0006 |
| TPXL-1 vs. SPD-2 | 14.25 | No | ns | 0.6655 |
| TPXL-1 vs. SPD-5 | -33.07 | Yes | ** | 0.0028 |
| TPXL-1 vs. TBG-1 | -4.446 | No | ns | 0.9652 |
| TPXL-1 vs. TAC-1 | -24.80 | No | ns | 0.1399 |
| TPXL-1 vs. PLK-1 | 49.61 | Yes | **** | <0.0001 |
| RSA-1 vs. RSA-2 | -9.346 | No | ns | 0.9002 |
| RSA-1 vs. SPD-2 | 41.71 | Yes | **** | <0.0001 |
| RSA-1 vs. SPD-5 | -5.608 | No | ns | 0.9652 |
| RSA-1 vs. TBG-1 | 23.02 | No | ns | 0.0790 |
| RSA-1 vs. TAC-1 | 2.661 | No | ns | 0.9652 |
| RSA-1 vs. PLK-1 | 77.07 | Yes | **** | <0.0001 |
| RSA-2 vs. SPD-2 | 51.06 | Yes | **** | <0.0001 |
| RSA-2 vs. SPD-5 | 3.739 | No | ns | 0.9652 |
| RSA-2 vs. TBG-1 | 32.36 | Yes | ** | 0.0014 |
| RSA-2 vs. TAC-1 | 12.01 | No | ns | 0.8399 |
| RSA-2 vs. PLK-1 | 86.42 | Yes | **** | <0.0001 |
| SPD-2 vs. SPD-5 | -47.32 | Yes | **** | <0.0001 |
| SPD-2 vs. TBG-1 | -18.70 | No | ns | 0.2609 |
| SPD-2 vs. TAC-1 | -39.05 | Yes | ** | 0.0014 |
| SPD-2 vs. PLK-1 | 35.36 | Yes | ** | 0.0020 |
| SPD-5 vs. TBG-1 | 28.62 | Yes | ** | 0.0061 |
| SPD-5 vs. TAC-1 | 8.269 | No | ns | 0.9307 |
| SPD-5 vs. PLK-1 | 82.68 | Yes | **** | <0.0001 |
| TBG-1 vs. TAC-1 | -20.35 | No | ns | 0.2689 |
| TBG-1 vs. PLK-1 | 54.06 | Yes | **** | <0.0001 |
| TAC-1 vs. PLK-1 | 74.41 | Yes | **** | <0.0001 |

## METHODS

### Contact for reagent and resource sharing

Further requests and information for resources and reagents should be directed to and will be fulfilled by the Lead Contact, Jeffrey Woodruff.

### Experimental model and subject details

*C. elegans* worm strains were grown on nematode growth media (NGM) plates at 16-23°C, following standard protocols (www.wormbook.org). Worm strains used in this study are listed in Table S1 and created using CRISPR (Paix et al., 2015; Paix et al., 2017), MosSCI (Frokjaer-Jensen et al., 2008), or microparticle bombardment. Cas9 enzyme was purified by the Protein Expression Facility at MPI-CBG. For expression of recombinant proteins, we used suspended SF9-ESF *S. frugiperda* insect cells grown at 27**°**C in ESF 921 Insect Cell Culture Medium, Protein-Free (Expression Systems), supplemented with Fetal Bovine Serum (2% final concentration).

### RNAi treatment

RNAi was done by feeding using *sur-6*, *gpr-2*, *csnk-*1, and *perm-*1 feeding clones from the Ahringer and Vidal collections (Source BioScience)(Rual et al., 2004). The *spd-5* feeding clone targets a region that is reencoded in our MosSCI transgenes (Woodruff et al., 2015). Bacteria were seeded onto nematode growth media (NGM) supplemented with 1 mM isopropyl *b*-D-1-thiogalactopyranoside (IPTG) and 100 µg mL^-1^ ampicillin. For *perm-1* feeding plates, 0.1 mM IPTG was used. L4 hermaphrodites were grown at 23°C for 24-28 hours for all conditions except for *perm-1*, which was at 20°C for 18-19 hours.

### Drug treatment of semi-permeable embryos

For all drug treatments, *C. elegans* embryos were permeabilized using *perm-1* RNAi (Carvalho et al., 2011) and dissected into a 62% solution of ESF-921 Media (Expression Systems). To arrest the embryos at metaphase, MG-132 (EMD Millipore) was used at 10 µM in 62% ESF, diluted from a 10 mM stock concentration in EtOH. To inhibit PLK-1, BI-2536 (Advanced ChemBlocks Inc.) was used at 10 µM, diluted from a 10 mM stock in ethanol. To inhibit PP2A, LB-100 (SelleckChem) was used at 10 µM, diluted from a 10 mM stock in dH_2_O. Nocodazole (Sigma) was diluted from a 5 mg/ml stock concentration in DMSO. Samples were flushed with water or M9 after each experiment to test for permeability (water will cause swelling and M9 will cause shrinking).

### Construction of the FLUCS Microscope

To measure the physical material state of centrosomes inside living *C. elegans* embryos, we performed intracellular flow perturbations by employing the previously published technology FLUCS (Mittasch et al., 2018). The FLUCS setup consisted of three major modules: (i) an infrared laser scanning unit for thermal manipulations, (ii) a microscope allowing to simultaneously induce thermal patterns and to perform high-sensitivity fluorescence imaging, and (iii) a heat management stage.

(i) The infrared laser scanning unit consists of a fiber-based infrared Raman laser (CRFL-20-1455-OM1, 20 Watts, near TEM00 mode profile, Keopsys, France) with a wavelength of 1455 *nm*, operated in continuous-wave mode and linearly polarized using a polarizing beam splitter cube (CCM1-PBS254, Thorlabs, USA). To precisely correct for the divergence of the laser beam, a telescope was used, composed of two telescope lenses with focal lengths of f_1=100 mm and f_2=150 mm (AC254-C series, Thorlabs, USA), respectively. A lambda-half plate (waveplate, 1/2 1550 Edmund optics, USA) was used to rotate the linearly polarized laser light to match the optical axis of the acoustic-optical deflector (AOD). A variable optical beam expander (4x expander, 36100, Edmund optics, USA) allows control of the beam diameter (∼1.5 mm beam diameter at back-focal-plane was used) without changing the size of the scan pattern. Rapid (up to 1 MHz update rate) and precise (down to 100 nm) infrared laser scanning was achieved by utilizing two-dimensional AOD (AA.DTSXY-A6-145, Pegasus Optik, Germany), electronic oscillators (AA.DRFAI0Y-B-0-x, Pegasus Optik, Germany), and electronic amplifiers 2.5 *W* (AA.AMPA-B-34-20.4, Pegasus Optik, Germany). The AOD was controlled by generating analog signals using a custom software in LabVIEW (National instruments, USA) in combination with a PCI controller card (PCIe 6369, National Instruments, USA). To precisely translate the AOD-induced beam scanning into the back-focal-plane of the microscope objective lens a telescope composed of two telescope lenses with focal lengths of f_3=f_4=300 mm (AC254-C series, Thorlabs, USA) was used. A dichroic mirror (F73-705, AHF, Germany) was used to couple the infrared laser beam into the light path of the microscope (IX83, Olympus, Japan), by selectively reflecting the infrared light but transmitting visible wavelengths which were used for fluorescence imaging.

(ii) The microscope was equipped with Brightfield (BF) and fluorescent imaging optics. For simultaneous high-resolution fluorescence imaging and precise infrared laser scanning, an infrared-coated microscope objective lens (60x UPLSAPO NA=1.2, W-IR coating, Olympus, Japan) was used, which was operated with heavy water (D2O) as immersion liquid to reduce undesired infrared laser light absorption in the immersion layer. For Brightfield illumination a high-power LED (M565L3, Thorlabs, USA) in combination with dedicated LED driver (LEDD1B, Thorlabs, USA) was used. For confocal fluorescence imaging, a VisiScope confocal imaging system (Visitron, Germany) coupled to a Yokogawa CSU-X1-A12 scan head and an iXON Ultra EMCCD camera (Andor, Ireland) were used.

(iii) The heat management unit consisted of a thin sample mounting chamber based on a standard cover slip (18 × 18 × 0.17 *mm*) (Menzel, Germany) facing the objective lens, and a high thermal conductive sapphire cover slide (thermal conductivity of 27.1 *W*⁄*m* · *K*, SMS-7521, UQG Optics, UK) closing the sandwich-like chamber from the top. To efficiently remove the induced heat from the samples, *e.g. C. elegans* embryos, the sapphire slide was actively cooled from room temperature to 17°*C*. This active cooling was performed by using Peltier elements (TES1-127021, TEC, Conrad) glued to the sapphire slides. The cooling power of the Peltier elements was controlled by a PID hardware controller (TEC-1089-SV, Meerstetter Engineering, Swiss). A custom-built water-cooling stage was used to dissipate the heat produced by the Peltier elements. The height of the buffer-filled chamber was defined using polystyrene beads (Polybead, Polysciences, Germany) with a diameter of 15 μ*m*. The height of the resulting chamber was measured by locating the upper and lower chamber surface using a piezo stage.

### Application of FLUCS within embryos

Late L4 hermaphrodites were grown for 18-19 hours on standard NGM or *perm-1* feeding plates. Worms were then dissected on an 18 mm x 18 mm coverslip (0.17 mm thickness) in 6 µL of M9 buffer or 62% ESF 921 (for permeabilized embryos) with 15 µm polystyrene beads. The sample was placed onto a sapphire microscope slide equipped with Peltier cooling elements, then the coverslip sealed with dental silicone (Picodent twinsil, Picodent, Germany). The cooling stage and sample were then mounted on the FLUCS microscope stage. Embryos were identified and staged using a 10x air objective, then imaged with a 60x 1.2 NA Plan Apochromat water immersion objective (Olympus) using 488 nm and 561 nm laser illumination, 1X1 binning, and 2s intervals.

Hydrodynamic flows were generated by scanning the 1455 nm laser through either 1) center of centrosome or 2) through the cytoplasm for velocity calibration. Custom-written LabVIEW software superimposes the scan path of the infrared laser with the high-resolution image of the camera. The sub-pixel alignment of the induced flow field and the camera image was verified routinely before the embryonic experiments by using fluorescent tracer particles immersed in a highly viscous sucrose solution. FLUCS experiments used unidirectional but repeated laser scans with 1.5 *kHz* scan frequency, a scan length of 30 μ*m*, and three different laser powers (25 mW, 32 mW, and 40 mW).

Centrosomes were targeted for FLUCS at metaphase, anaphase, or telophase. Centrosomes were affected by FLUCS between 30-60 s. For experiments requiring drug treatment, worms were dissected in 6 µL of the specific drug solution and quickly placed on the microscope within 1-2 minutes. To maintain consistency of drug treatment duration, only embryos found exactly at prometaphase (for metaphase experiments) and metaphase (for anaphase experiments) were then targeted for FLUCS. Temperature-sensitive worms were dissected in cold 62% ESF-921 media on a cooled dissecting scope and quickly mounted onto the cooling stage, which was maintained at 17°C. At prometaphase, temperature was upshifted to 25°C for 1 minute, then decreased to 17°C. Centrosomes were then targeted for FLUCS at metaphase.

### Confocal microscopy and live-cell imaging

Adult worms were dissected in M9 before being mounted on a 5% agar pad for imaging. For live cell imaging with drug treatments, *perm-1* adult worms were dissected in 8-10 µL of 62% ESF 921 with 15 µm polystyrene beads (Sigma-Aldrich) on a 22 x 50 mm coverslip. Samples were mounted on a 1 mm thick glass slide with 2 x 6 mm laser cut holes 30 mm apart (Potomac), to produce a flow chamber. In one open chamber, 40 µL of the drug solution in 62% ESF was added during prometaphase. Liquid was wicked from the opposite chamber using a Kimwipe to then allow more of the drug solution to be added to the sample. To arrest embryos in metaphase, *perm-1* adult worms were dissected in 10 μM MG-132 solution. Cell cycle stage was indicated based on mCherry::HIS-58 fluorescence and cell morphology (metaphase = aligned chromosomes; anaphase = chromosomes separate; telophase = chromosomes de-condense and cytokinetic furrow ingresses).

Time-lapse images were taken using an inverted Nikon Eclipse Ti microscope with a Yokogawa spinning disk confocal head (CSU-X1), piezo Z stage, and an iXon Ultra EMCCD camera (Andor), controlled by Metamorph software. On this system, the 60x 1.4 NA Apochromat oil objective was used to acquire 36 x 0.5 µm Z-stacks every 10 seconds with 100 ms exposures and 2×2 binning. For PCM localization in *csnk-1(RNAi)* embryos, and PP2A localization, time-lapse images were acquired with an inverted Nikon Eclipse Ti2-E microscope with a Yokogawa confocal scanner unit (CSU-W1), piezo Z stage, and an iXon Ultra 888 EMCCD camera (Andor), controlled by Nikon Elements software. For most experiments, we used a 60x 1.2 NA Plan Apochromat water immersion objective to acquire 35 x 0.5 µm Z-stacks every 10 seconds with 100 ms exposures and 2×2 binning. Simultaneous imaging with the 488 nm and 561 nm lasers was achieved using an OptoSplit II beam splitter (Cairn). For LET-92::GFP imaging, a 100x 1.35 NA Plan Apochromat silicone oil objective was used to acquire 11 x 0.5 µm Z-stacks in 20 second intervals with 100 ms exposures and 2×2 binning. Images in Figure 3 were taken using an inverted Olympus IX81 microscope with a Yokogawa spinning-disk confocal head (CSU-X1), a 60x 1.2 NA Plan Apochromat water objective, and an iXon EM + DU-897 BV back illuminated EMCCD (Andor).

### *spd-2(or188ts)* temperature shift assay

JWW69 (control) and JWW89 (*spd-2(or188ts)*) strains were used for imaging. Sequencing of JWW89 confirmed a single point mutation in *spd-2* resulting in a glycine to glutamic acid amino acid substitution (G615E) as described in Kemp et. al. Both worm strains propagated at 16°C, which is the permissive temperature for *spd-2(or188ts)*. To prepare the embryos for imaging, a metal block was buried halfway in wet ice. A 24×60 mm glass coverslip (thickness of 1) and a flow chamber slide were placed over the cold block. To prevent sticking of the glass to the cold block due to water condensation, two Kimwipes were placed between the glass and the cold block. To minimize exposure to elevated temperatures during embryo dissection, the glass stage on the dissecting microscope stand was placed in a 4°C fridge and left to cool for approximately 10 min. Once everything was cold, 10µL of cold M9 plus 15 µm polystyrene (Sigma) beads was pipetted to the middle of the 24×60 mm cover slip.

For each worm strain, plates were transported inside the ice bucket directly contacting ice to the dissecting microscope area. The microscope glass stage was taken out from the fridge and assembled into its place. Three to four adult worms containing a single row of eggs were transferred to the M9 plus beads on the cover slip still located on top of the cold block. The coverslip was transferred to the dissecting scope and the worms cut open using 22G needles. The coverslip was mounted on the flow chamber slide, then the edges of the cover slip were sealed using clear nail polish. The sample was moved to the imaging room on the cold block.

The Nikon Eclipse Ti2 microscope described above was used for imaging. Embryos were staged using a 10X air objective, then imaged with a 60X NA 1.2 water objective. To rapidly raise the temperature of the sample (up-shift), 40 µl of 25°C M9 was pipetted into the flow chamber well. 30 x 0.5 µm Z stacks were collected every 10 s using simultaneous illumination with 488 nm and 561 nm lasers (14.7% and 17.7% intensity respectively), 2×2 binning, 100 ms exposures.

### Protein expression and purification

All expression plasmids are listed in Table S2. SPD-5, SPD-2, and PLK-1 proteins were expressed using the FlexiBAC baculovirus system (Lemaitre et al., 2019) and purified as previously described (Woodruff and Hyman, 2015; Woodruff et al., 2015), with the following exception: SPD-2 was stored in its uncleaved form (MBP-TEV-SPD-2).

### *In vitro* SPD-5 condensate disassembly assay

SPD-5 condensates were formed by diluting 10 μM SPD-5 (1:10 mixture of SPD-5 and SPD-5::TagRFP) in Condensate buffer (25 mM HEPES, pH 7.4, 150 mM KCl) containing polyethylene glycol 3350 (Sigma) and fresh 0.5 mM DTT. Before use, the SPD-5 stock solution was centrifuged for 5 min at 80,000 rpm to remove residual aggregates. 5 min after formation, SPD-5 condensates were placed in glass-bottom 96-well dishes (Corning, 4850, high content imaging dish) pre-cleaned with 2% Hellmanex and washed in water. For each sample, half was placed in the well undisturbed (control), and the other half was diluted 10-fold, pipetted 5 times, then placed in a well (induced disassembly). 96-well plates were imaged on an inverted Nikon Ti-E microscope using a 60x NA 1.4 Plan Apochromat oil objective, a Zyla cMOS camera (Andor), and MicroManager control software. For each image, SPD-5 condensates were identified through applying a threshold then using the particle analyzer function in FIJI. When analyzing condensate formation, we report the sum of the integrated intensities of each condensate per image (total condensate mass). Survival % plotted in Figure 5 assumes a 10-fold loss in total condensate mass due to dilution.

### Quantification and statistical analysis

Images were analyzed with FIJI (https://fiji.sc/), R (https://www.r-project.org/), and GraphPad Prism (https://www.graphpad.com). For FLUCS experiments, centrosome deformation was calculated by measuring the long axis (orthogonal to the flow direction) of PCM-localized SPD-5 at the initial and final time points of PCM deformation, prior to fracture (defined below). The deformation rate equaled the difference in PCM lengths divided by the time interval. Centrosome fracture was measured using line scans across the long axis of PCM-localized SPD-5. Fracture was scored if signal dropped to cytoplasmic levels over three consecutive pixels on the long axis across the entire flow path, and if this signal gap persisted for the rest of the images. For all other experiments, centrosome tracking and measurement was conducted using max intensity projections, correction for photobleaching, followed by thresholding and particle analysis. Thresholds were determined using: mean background intensity of the cytoplasm + b*(standard deviation of background), where b represents an integer value that is identical for all samples within an experiment. The integrated fluorescence density for the centrosome-localized signals were normalized to either the first intensity value or max intensity value (Figure 3) and plotted over time.

All data are expressed as the mean ± 95% confidence intervals as stated in the figure legends and results. The value of n and what n represents (e.g., number of images, condensates or experimental replicates) is stated in figure legends and results. Normality tests were first performed before applying statistical tests. Statistical tests were performed with GraphPad Prism.

